# Inhibition of nitric oxide synthase transforms carotid occlusion-mediated benign oligemia into *de novo* large cerebral infarction

**DOI:** 10.1101/2024.07.18.604214

**Authors:** Ha Kim, Jinyong Chung, Jeong Wook Kang, Dawid Schellingerhout, Soo Ji Lee, Hee Jeong Jang, Inyeong Park, Taesu Kim, Dong-Seok Gwak, Ji Sung Lee, Sung-Ha Hong, Kang-Hoon Je, Hee-Joon Bae, Joohon Sung, Eng H. Lo, James Faber, Cenk Ayata, Dong-Eog Kim

**Author notes:** **Address correspondence to:** Dong-Eog Kim, National Priority Research Center for Stroke, Dongguk University Ilsan Hospital, 27, Dongguk-ro, Ilsandong-gu, Goyang 10326, Republic of Korea. **Authorship note:** HK and JC are co-first authors and contributed equally to this work.

## Abstract

It remains unclear why unilateral proximal carotid artery occlusion (UCAO) causes benign oligemia, without progressing to cerebral infarction, in mice, yet leads to a wide variety of outcomes (ranging from asymptomatic to death) in humans. We hypothesized that inhibition of NOS both transforms UCAO-mediated oligemia into full infarction and expands pre-existing infarction. In support, intraperitoneal administration of Nω-nitro-L-arginine methyl ester (L-NAME) followed by UCAO induced large-arterial infarction in mice, unlike UCAO alone. Six-hour laser-speckle-contrast imaging detected spreading ischemia in mice with infarction as assessed at 24h. In agreement with vasoconstriction/microthrombus formation shown by intravital microscopy, the NO-donor, molsidomine and the endothelial-NOS- activating antiplatelet, cilostazol, attenuated or prevented progression to infarction. Moreover, UCAO without L-NAME caused infarction in mice with hyperglycemia and hyperlipidemia, which, in turn, were associated with greater symmetric dimethylarginine (SDMA) levels. Further, increased levels of glucose and cholesterol associated with significantly larger infarct volumes in 438 consecutive patients with UCAO-mediated infarction. Lastly, Mendelian randomization identified a causative role of NOS inhibition, particularly in elevated SDMA concentration, in ischemic stroke risk. Therefore, NOS activity is a key factor determining the fate of hypoperfused brain following acute carotid occlusion, where SDMA could be a potential risk predictor.

## Introduction

Carotid artery occlusion (CAO) accounts for up to ∼15% of ischemic stroke (1), which is a leading cause of death and disability (2). As vascular imaging becomes more accessible, carotid occlusive diseases are diagnosed more frequently. CAO can lead to a wide variety of outcomes, ranging from death to severe stroke-related impairment to asymptomatic (3), and the mechanisms behind this variability remain incompletely understood.

Cerebral infarction after proximal arterial occlusions is uncommon, unless there is blood flow reduction below a critical threshold, artery-to-artery embolism, or tandem occlusion, because cerebral circulation is highly collateralized (4–6). Although anatomic collateral abundance varies among individuals, adequate collateral blood flow cannot be solely attributed to anatomy. Vascular reactivity within and downstream of the collateral network in the setting of large artery obstruction likely plays a pivotal role in adequate collateral compensation (7). Cerebral vasoreactivity is affected by vascular risk factors that can impair endothelial function including, notably, the modulation of NO-dependent vasodilation and regulation of interactions with platelets and leukocytes (8–10). Many studies have shown that NO deficiency and NOS inhibition contribute to lesion size in acute cerebral infarction (11–13). However, to the best of our knowledge, no studies have investigated if NOS inhibition-mediated NO deficiency, in the absence of preexisting acute cerebral infarction, can cause acute infarction in the oligemic but non-infarcted hemisphere.

Recently, NO deficiency in mice was found to impair cerebral microcirculation adaptation to unilateral common carotid artery occlusion (UCAO) (14). During stroke induction by filament occlusion of the proximal middle cerebral artery (MCA), a branch of the internal carotid artery which itself is a branch of the common carotid artery, cerebral blood flow (CBF) needs to decrease to ∼20% of baseline to successfully trigger cerebral infarction (15). Despite strain-specific differences in pial and circle of Willis collateral status, acute UCAO usually reduces ipsilateral CBF to ∼60% of baseline (16), and this level of CBF reduction (i.e., benign oligemia) is insufficient to cause cerebral infarction (17).

Here, we tested our hypothesis that UCAO-mediated hypoperfusion that is insufficient to cause ischemic brain damage can lead to cerebral infarction in the presence of NO deficiency by a single-dose NOS inhibitor (NOSi): Nω-nitro-L-arginine methyl ester (L-NAME). We corroborated the mouse experiments (n=900) with a clinical study of 438 consecutive UCAO- stroke patients and a Mendelian randomization study on endogenous NOSi (dimethylarginines), thereby investigating the presence of a causative link between NOS inhibition and risk of ischemic stroke in humans. We also demonstrate that combining UCAO with NOS inhibition could serve as a unique *in vivo* stroke model for preclinical research, such as stroke occurrence investigations that are not achievable with present approaches.

## Results

### UCAO induced cerebral infarction in ∼75% of C57BL/6 and BALB/c mice pre-treated with L-NAME

UCAO alone rarely induced infarction in C57BL/6 mice (2%, 1/47): only one of 37 (3%) subjected to UCAO for 24 h had an infarct, which was small, and none of 10 (0%) exposed to UCAO for 7 d had infarcts (Figure 1A). As expected, neither L-NAME alone (n=12) nor sham surgery (n=7) led to infarction (Figure 1A). In contrast, UCAO combined with prior administration of a single intraperitoneal dose of L-NAME (L-NAME+UCAO) caused infarction in most animals (74%, 55/74; Figure 1, A and B), as shown by 2,3,5- Triphenyltetrazolium chloride (TTC) staining performed after premature death (lethal infarction) or pre-planned sacrifice on day 1, 2, 3, or 6. Most infarcted mice had large lesions (>100 mm^3^, 77%, 42/55), as quantified after death (100%, 23/23) or sacrifice (59%, 19/32). Mean±standard error (SE) infarct volumes of the surviving mice were 115±20, 74±30, 55±25, and 9±9 mm^3^ at the above time-points respectively. This declining trend reflects that mice with larger infarcts are more likely to die before planned euthanasia. L-NAME+UCAO-mediated severe (large or lethal) infarction occurred more frequently in BALB/c mice, whereas infarction occurred less often in SV129 mice. For further details, see Supplemental Material. Of female C57BL/6 mice receiving L-NAME+UCAO, 68% (15/22) had infarction by day 1, a result which did not differ significantly from the aforementioned male data: 69% (29/42), *P*=0.94. Approximately half of infarcted females had large lesions (53%, 8/15) when assessed after death (100%, 2/2) or sacrifice (46%, 6/13).

**Figure 1.**
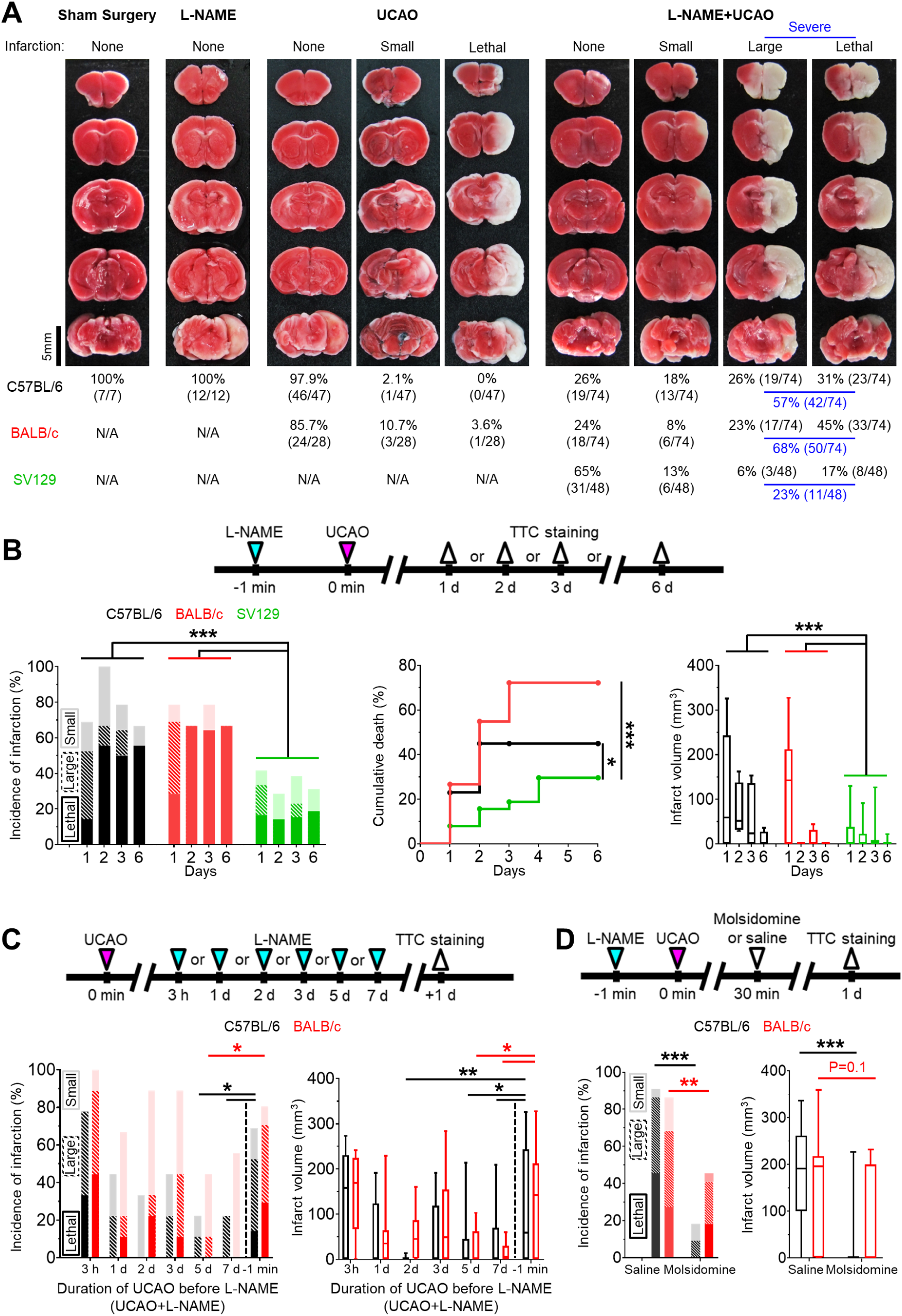
L-NAME+UCAO in mice can induce acute large cerebral infarction, which could be rescued by the NO donor molsidomine. (**A**) TTC staining of the brains from a representative C57BL/6 mouse (except for UCAO - lethal infarction of a BALB/c mouse) with no, small (≤100 mm^3^), or large (>100 mm^3^) infarction in each of the four (sham surgery, L- NAME, UCAO, and L-NAME+UCAO) groups. (**B**) Top, experimental timeline for L- NAME+UCAO. Bottom-left, incidence of cerebral infarction by a pre-specified sacrifice time- point. Light, striated, and dark colors represent small, large, and lethal infarction, respectively. Bottom-middle, stroke-related mortality curves. Bottom-right, infarct volumes of the mice that survived until the pre-specified sacrifice time-points. (**C**) Top, experimental timeline for L- NAME post-treatment (UCAO+L-NAME): L-NAME administration at a different post-UCAO time-point, followed by sacrifice 24 h later. Bottom-left, incidence of cerebral infarction. Bottom-right, infarct volumes of the mice that survived until the 24 h sacrifice time-point. For easier comparison with the corresponding L-NAME pre-treatment (L-NAME+UCAO) data in (**B**), the 1 d infarct-incidence and -volume bars are presented here again (−1 min). (**D**) Top, timeline for molsidomine experiments. Bottom-left, incidence of cerebral infarction by 24 h sacrifice time-point in mice treated with a single intraperitoneal dose of molsidomine or saline. Bottom-right, infarct volumes of the surviving mice. Infarct volumes are presented as box-and-whiskers plots. \**P*<0.05, \*\**P*<0.01, and \*\*\**P*<0.001. N/A, not applicable.

### Administering L-NAME after, rather than before, prolonged UCAO resulted in lower incidence of infarction and smaller lesion size

Prolonged (5 or 7 d) UCAO, but not shorter (3 h, 1–3 d) UCAO, before L-NAME administration (UCAO+L-NAME) in C57BL/6 and BALB/c mice (n=9/time-point/strain) induced infarction (∼50%) less frequently and produced (∼70%) smaller lesions as measured 24 h after L-NAME administration (Figure 1C), when compared to the aforementioned L-NAME+UCAO (for the 1 d sacrifice groups). For further details, see Supplemental Material.

### NO-donor, molsidomine, prevented or attenuated L-NAME+UCAO-induced infarction

As shown in Figure 1D, infarction (by 1 d, including lethal infarction) occurred less frequently in molsidomine-treated mice (C57BL/6: 18%, 4/22; BALB/c: 46%, 10/22) than in saline-treated mice (C57BL/6: 91%, 20/22; BALB/c: 86%, 19/22); both *P* were less than 0.01 (Fisher’s exact test). None of the molsidomine-treated C57BL/6 mice (0%, 0/22) died, whereas 10/22 (46%) of saline-treated mice died (*P*<0.001, Fisher’s exact test). Four (18%) of the molsidomine- treated BALB/c mice (n=22) died, whereas six (27%) of the saline-treated mice (n=22) died (*P*=0.72, Fisher’s exact test). Mean±SE infarct volumes of the C57BL/6 mice that survived until sacrifice were significantly lower after molsidomine vs. saline treatment (19±13 mm^3^ vs. 177±32 mm^3^; *P*<0.001, two-sample *t* test). In BALB/c mice, there was a trend toward molsidomine treatment-related reduction in infarct volume (63±25 mm^3^ vs. 130±30 mm^3^; *P*=0.10, two-sample *t* test).

### L-NAME+UCAO induced infarction despite cortical blood flow initially being as high as ***∼65% in the core region***

In C57BL/6 mice (n=19) and BALB/c mice (n=24), laser speckle contrast imaging (LSCI) was performed at baseline and for 6 h after L-NAME+UCAO (Figure 2A). At 0-10 min after L-NAME+UCAO, both mouse strains had similarly high mean±SE regional cortical blood flow (rCoBF) in the oligemic core: 68±2% and 64±3%, respectively, of their pre-intervention baseline values in the core region (region of interest [ROI]-1, Supplemental Figure 1). Notably, mean rCoBF values at 0-10 min were higher than 30% in every cortical ROI of all mice. Although cerebral infarction and stroke-related death are highly unlikely outcomes after oligemia or non-critical reduction of rCoBF at this level, they occurred in more than half the animals (61%, 27/44): specifically, 53% (10/19, including one case of lethal infarction) of C57BL/6 and 68% (17/25, including six cases of lethal infarction) of BALB/c mice. For further details, see Supplemental Material.

**Figure 2.**
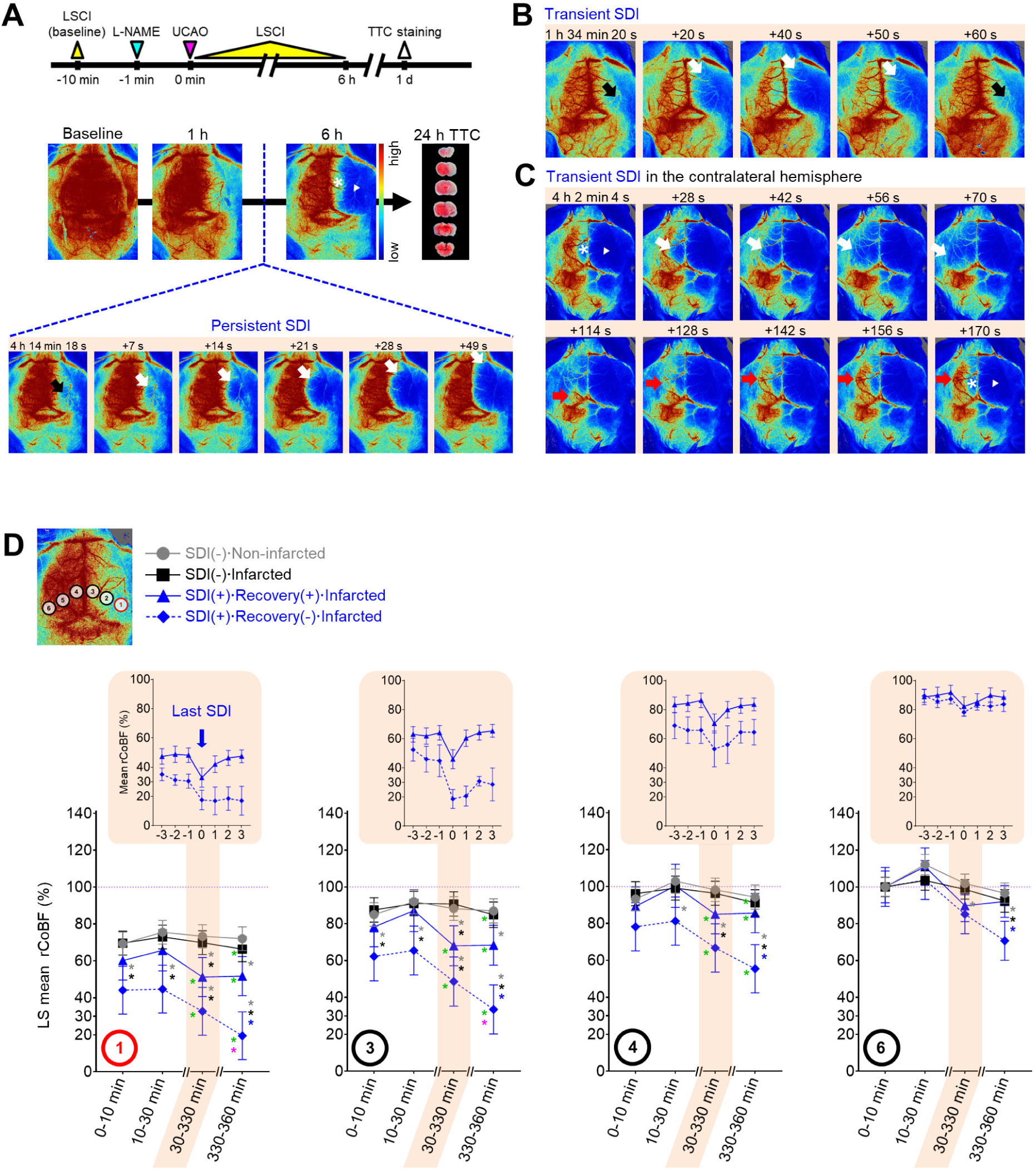
LSCI for 6 h after L-NAME+UCAO detected SDI in ∼40% of mice with cerebral infarction assessed at 24 h, but in none without. (**A**) Timeline for experiments and persistent SDI in the hemisphere ipsilateral to UCAO in a BALB/c mouse. SDI began, with a substantial drop in rCoBF from the oligemia (>50%, black arrow) to the severe ischemia level (<30%, white arrows), in the core region (ROI-1), spreading anteromedially to encompass the majority of the ipsilateral hemisphere. Except for the medial portion (*), there was no rCoBF recovery (white arrow-head). See Supplemental Video 1. (**B**) Transient SDI (black and white arrows) in the ipsilateral hemisphere of another BALB/c mouse. See Supplemental Video 2. (**C**) Transient SDI with CoBF recovery (white and red arrows) in the contralateral hemisphere of another BALB/c mouse. See Supplemental Video 3. (**D**) LS mean rCoBFs with 95% CIs, calculated after stratifications by the occurrence of SDI (with (+) or without (-) rCoBF recovery up to 6 h) and cerebral infarction (up to 24 h). Gray, black, and blue * indicate *P*<0.05 vs. the SDI(-)ꞏNon-infarcted group, SDI(-)ꞏInfarcted group, and SDI(+)ꞏRecovery(+)ꞏInfarcted group, respectively, at each time period. Green and pink * indicate *P*<0.05 vs. 10-30 and 30-330 min, respectively. Additionally, rCoBF values (mean±SE) from 3 min before to 3 min after the last SDI are displayed for each SDI-positive group (upper graphs in the shaded areas).

### Spreading ischemia identified mice that progressed to infarction after L-NAME+UCAO

LSCI for 6 h after L-NAME+UCAO detected spreading ischemia (known to be linked to spreading depolarization [SDI]) (18, 19) in ∼40% of mice with infarction, assessed at 24 h, but in none without. During the 6 h LSCI monitoring, single or recurrent bouts of spontaneous (and prominent) SDI were observed (Figure 2 and Supplemental Videos 1-3) in 40% (10/25; 4 and 6, respectively) of mice that had cerebral infarcts by 24 h (three C57BL/6 mice and seven BALB/c mice including two with lethal infarction). In these 10 animals, mean±SE (median, IQR) rCoBF in the core at 0-10 min after L-NAME+UAO was 53.7±4.5 (52.7, 41.2-64.1) %. Their first SDIs occurred at 127±33 (69, 58–254) min, and pre-SDI rCoBF (at 3 min before the first SDI) was 44.6±6.2 (41.5, 34.2-57.5) %. Note that SDI was not observed in any of the 17 mice without infarcts (*P*=0.004, Fisher’s exact test). Moreover, severe (i.e., large or lethal) infarction occurred in all SDI-positive mice except for one C57BL/6 mouse that had a small infarct. SDI usually began in the posterolateral region of the hemisphere ipsilateral to UCAO (ROI-1 or nearby; Figure 2), which was not the case in some mice. rCoBF (mean of the lowest values from individual SDI events) acutely dropped to less than 30% of the baseline in the ROI-1 of most (7/10) mice and to about 36-52% (36.5, 50.2, and 52.2%) in the remaining three mice (Supplemental Figure 2). Thus, these 10 animals’ mean±SE rCoBF value was 26.7±5.2%. The initial ischemic foci gradually expanded, usually toward the anteromedial portion of the ipsilateral hemisphere, thereby encompassing most of the ipsilateral hemispheric cortex. For further details, see Supplemental Material.

### L-NAME+UCAO-mediated serial changes in rCoBF differed depending on the occurrence of SDI (with or without rCoBF recovery up to 6 h) and infarction (up to 24 h)

In the core region (ROI-1), initially (0-10 and 10-30 min after L-NAME+UCAO), every group’s least squares (LS) mean rCoBF was higher than 30%, although the SDI(+)**·**Recovery(-)**·**infarcted group had a significantly lower LS mean value (∼50%) than the two SDI(-) groups (∼75%; Figure 2D). At 30-330 min, both SDI(+) groups had significantly lower LS mean rCoBF values than the two SDI(-) groups. In each SDI, as demonstrated by the upper graphs in the shaded areas of Figure 2D, which shows the last SDI data, rCoBF dropped to about 30% or lower in both SDI(+) groups, with or without subsequent rCoBF recovery, respectively. Thus, post-SDI LS mean (95% CI) rCoBF at 330-360 min was significantly lower in the SDI(+)**·**Recovery(-)**·**Infarcted group than in the SDI(+)**·**Recovery(+)**·**Infarcted group: 19.4% (6.5-32.4) and 51.7% (41.2-62.3), respectively. At 330-360 min, contrasting 10-30 min, all three infarcted groups had significantly lower LS mean rCoBF values. When compared with 30-330 min, the SDI(+)**·**Recovery(-)**·**Infarcted group only had a significantly lower LS mean rCoBF value at 330-360 min. Within each group, serial rCoBF changes were similar between C57BL/6 and BALB/c mice (Supplemental Figure 3 and Supplemental Material). As the ROIs were farther away from the core region (i.e., from ROI-1 to -6), LS mean rCoBF was higher, with fewer or no serial changes or inter-group differences (Figure 2D). For further details, see Supplemental Material and Supplemental Figure 4.

### Infarction following L-NAME+UCAO was not associated with systemic hypotension

Considering that combining UCAO with hypotension can induce focal ischemia (20), we monitored heart rate and BP as well as CoBF in order to investigate if L-NAME administration and/or UCAO associates with hemodynamic instability (Figure 3). Heart rate and systemic BP were not meaningfully different between groups, making it unlikely that systemic hypoperfusion causes SDI and infarction following L-NAME+UCAO. For further details, see Supplemental Material and Supplemental Figure 5 and 6.

**Figure 3.**
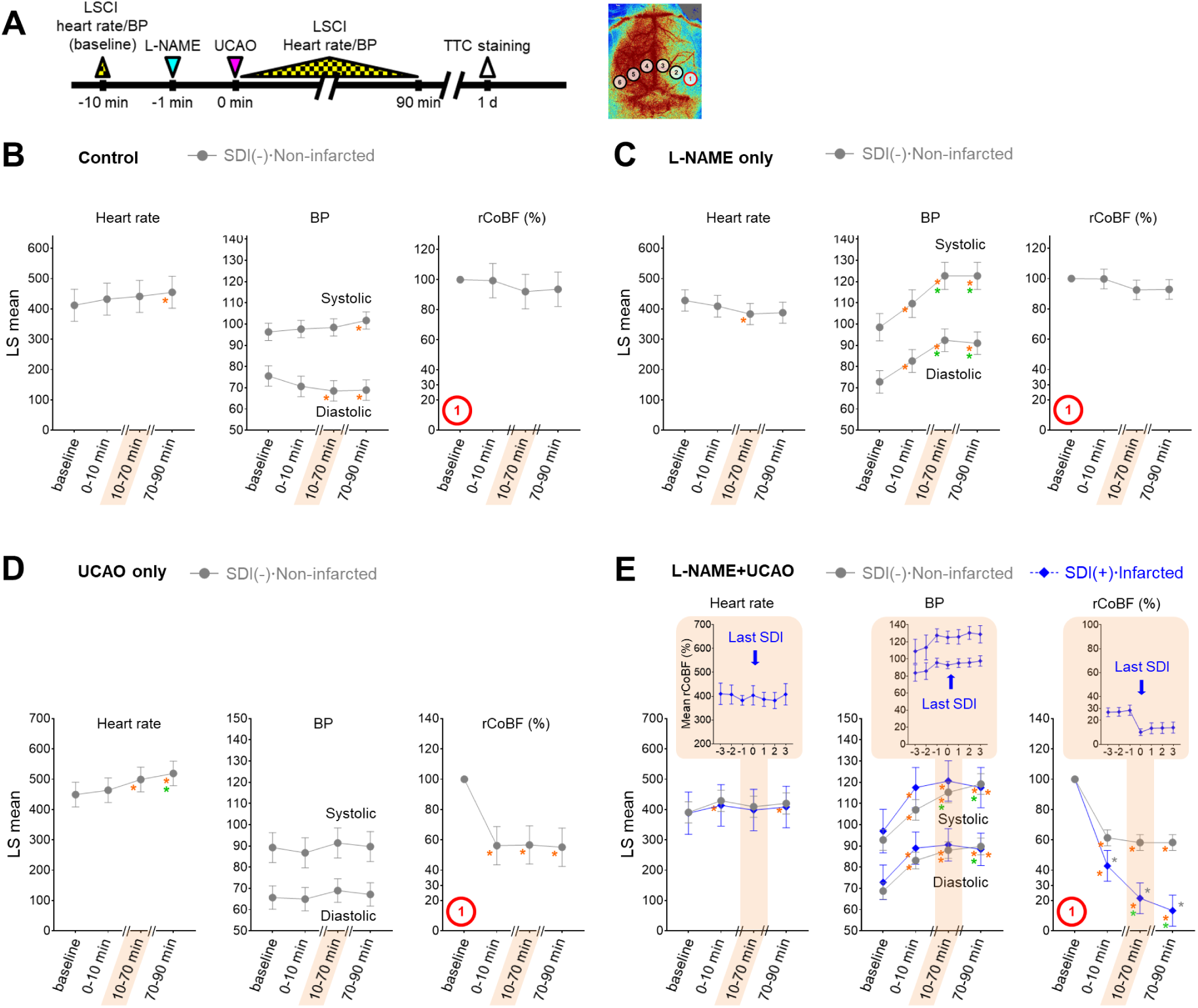
L-NAME+UCAO-mediated SDI and infarction were not associated with systemic hemodynamic instability. (**A**) Timeline for LSCI and heart rate/BP experiment. (**B**-**E**) Heart rate, systolic and diastolic BPs, and rCoBF (all as mean±SE) in the oligemic core (ROI-1): (**B**) saline-injected control group, (**C**) L-NAME only group, (**D**) UCAO only group, and (**E**) L-NAME+UCAO group, with stratifications by the occurrence of SDI (with (+) or without (-) rCoBF recovery up to 90 min) and cerebral infarction (up to 24 h). LS mean values with 95% CIs are presented in the graphs. Gray * indicates *P*<0.05 vs. the SDI(-)ꞏNon-infarcted group at each time period. Orange and green * indicate *P*<0.05 vs. baseline and 0-10 min. In addition to the statistical analyses, rCoBF values for the last SDI during the 10-70 min period are displayed for each SDI-positive group (as group mean±SE) in order to show SDI-related rCoBF drop to a trough level (inset graph in the shaded area). Corresponding heart rate and BP data (at the time-point of the lowest rCoBF) are also presented in inset graphs.

### Arteriolar constriction and microvascular thrombosis were associated with L-NAME+UCAO-mediated infarction, which the vasodilatory and anti-thrombotic drug cilostazol could prevent or attenuate

In thrombosis-related experiments (Figure 4), *in vivo* high-resolution microCT imaging to directly visualize thrombus using fibrin-targeted gold nanoparticles (Figure 4A) serially for 6 d after L-NAME+UCAO (Figure 4B) did not show evidence of large arterial thrombosis in C57BL/6 (n=14) and BALB/c (n=9) mice. H&E staining of brain tissue sections that were obtained 24 h after L-NAME+UCAO (5 C57BL/6 mice) showed arteriolar and capillary microthrombi, particularly in the infarcted hemisphere ipsilateral to UCAO (Figure 4C). Moreover, intravital two-photon microscopy imaging through a cranial window (Figure 5) into ipsilateral brain regions (serially by ∼4 or ∼24 h after L-NAME+UCAO in three C57BL/6 and 13 BALB/c mice) demonstrated, in real time, microvascular thrombosis (Figure 5C) as well as arteriolar constriction (Figure 5, A and B, and Supplemental Figure 7 and Supplemental Video 4). These impeded microcirculations and significantly reduced cortical vascular diameter and density (Supplemental Figure 8), particularly in mice with infarcts assessed by TTC staining at 4 or 24 h (Also see Supplemental Figure 9 for a representative non-infarcted mouse). Additionally, in line with the aforementioned mean onset time of the first SDI (∼2 h), platelet–leukocyte rolling and adhesion tended to be observed more often at ∼4 h rather than at ∼1 h (Supplemental Videos 5 and 6). Next, we tested whether L-NAME+UCAO-mediated infarction in C57BL/6 mice can be prevented or attenuated by pre-treatment with widely used antiplatelet drugs including aspirin, clopidogrel, and especially cilostazol, which is a vasodilatory antiplatelet drug that could increase endothelial NOS (eNOS) activity and NO production (21, 22). As Figure 4D shows, stroke-related mortality (by 6 d) was significantly lower in the cilostazol group than in the saline group (*P*=0.018, log-rank test): 50% (12/24) vs. 88% (21/24) by 6 d. The mortality tended to be lower in the clopidogrel group (67%, 16/24) and the aspirin group (71%, 17/24), compared to the saline group (*P*=0.074 and 0.077, respectively). The incidence of severe (i.e., large or lethal) infarction appeared to be lowest in the cilostazol group (63%, 15/24), followed by the clopidogrel group (71%, 17/24), and the aspirin group (83%, 20/24), and highest in the saline group (88%, 21/24): *P*=0.046, chi-square test between the cilostazol and saline groups. Additionally, mice that survived until 6 d did not exhibit significant inter-group differences in infarct volumes (two-sample *t* tests).

**Figure 4.**
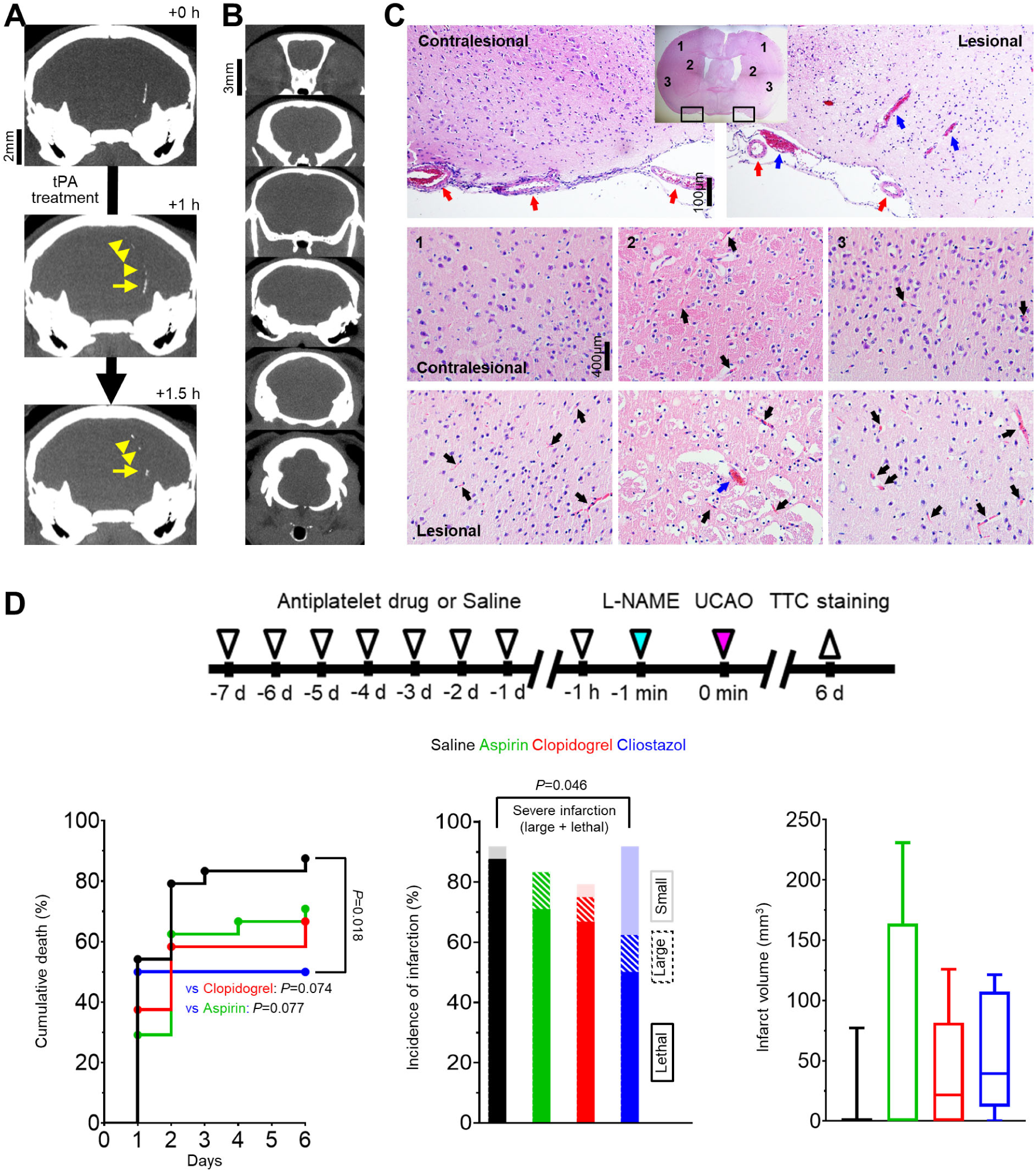
Abundant microthrombi were observed in infarcted mouse brain tissue 24 h post L-NAME+UCAO, corroborated by subsequent mouse experiments in which the vasodilatory antiplatelet cilostazol reduced the occurrence of severe infarction, probably, in part, via its antithrombotic effects. (**A**) Representative high-resolution *in vivo* microCT images showing that fibrin-targeted gold nanoparticles can clearly identify sizeable thrombi and their evolution (yellow arrows and arrow-heads) after tPA therapy for embolic stroke. (**B** and **C**) Assessment of the presence of macrovascular and microvascular thrombi: (**B**) no evidence of macrovascular thrombosis on microCT imaging (serially for 6 d) vs. (**C**) histologic (H&E staining) evidence of abundant microvascular thrombosis (black arrows) predominantly in infarcted brain tissue (sacrificed with cardiac perfusion). Venous thrombi are also apparent (blue arrows), whereas large arterial thrombi are not (red arrows). (**D**) Top, timeline for experiments to test if 8 d pre-treatment with an antiplatelet drug can show protective effects against L-NAME+UCAO-mediated infarction in C57BL/6 mice. Bottom-left, stroke-related mortality curves for the saline, aspirin, clopidogrel, and cilostazol groups. The mortality is lowest in the cilostazol group (*P*=0.018 vs. saline group, log-rank test with post-hoc pairwise comparisons). Bottom-middle, infarction incidence by 6 d. Light, striated, and dark colors represent small (≤100 mm^3^), large (>100 mm^3^), and lethal infarction, respectively. Severe (large or lethal) infarction was significantly less frequent in the cilostazol group (*P*=0.046 vs. saline group, chi-square test). Bottom-right, infarct volumes (box-and-whiskers plots) of the mice that survived until 6 d.

**Figure 5.**
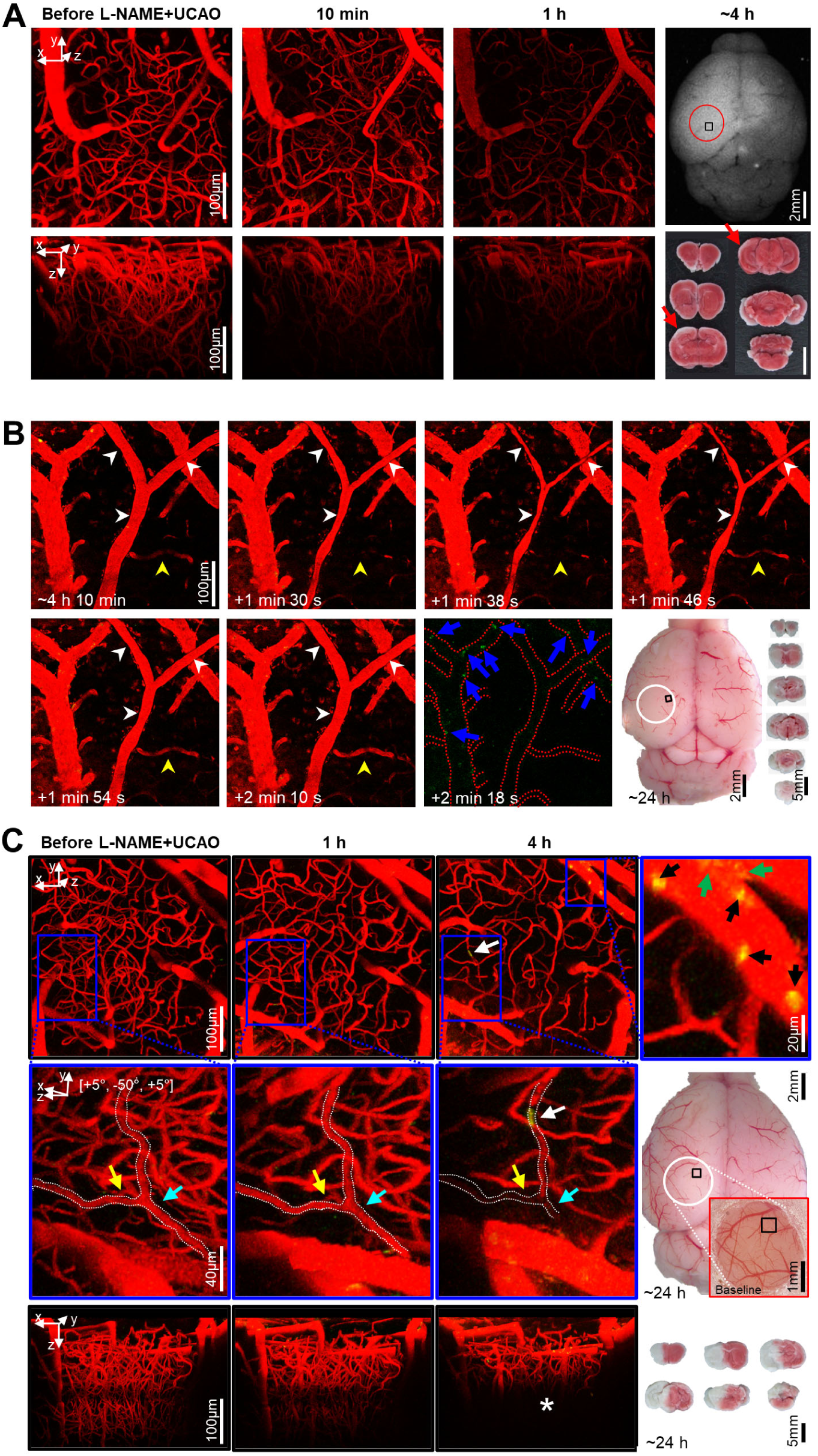
Intravital microscopy through a cranial window showed microvascular constriction and thromboembolism to impede distal microcirculation after L- NAME+UCAO. (**A**-**C**) Intravital images for the squared regions within the circles (far-right). (**A**) Stack images before and after L-NAME+UCAO in a BALB/c mouse. At 1 h, Texas-red-Dextran-positive vascularity is markedly decreased. At 4 h, post-mortem autofluorescence imaging and TTC staining (far-right) reveal whitish ischemic lesions (red arrows). (**B**) Transient constriction of the Y-shape arteriole in real-time (white arrow-heads) ∼4 h after L- NAME+UCAO in another BALB/c mouse (Supplemental Video 4; See also Supplemental Figure 7 for a different mouse case). Also note the congruent drop and rise in the blood flow of an adjacent arteriole (yellow arrow-heads), and Rhodamine-6G-positive platelets and leukocytes in the nearby venules (blue arrows). Post-mortem evidence of hemispheric infarction (bottom-row, far-right) suggests additional instances of vasoconstriction by ∼24 h. (**C**) Stack images before and after L-NAME+UCAO in another BALB/c mouse. At 4 h, Rhodamine-6G-positive platelets and leukocytes (green and black arrows) are abundant in arterioles (top-row, far-right). Moreover, the magnified views (middle-row) of the blue squared regions (top-row) show that capillary plugging (white arrows) accompanies distal flow reduction (cyan and yellow arrows). Also note decreased vascular density at 4 h (* in the bottom-row). See Supplemental Figure 8 for grouped quantification data for vascular diameter and density in all animals (n=16).

### UCAO, alone, without L-NAME administration, induced infarction in mice with hyperglycemia and hyperlipidemia, which in turn increased blood levels of symmetric dimethylarginine (SDMA, an endogenous NOSi)

Hyperglycemia and hyperlipidemia have been linked to NOS dysfunction in humans (23, 24). Therefore, we asked whether UCAO also induces infarction in mice after streptozotocin (STZ)-induced hyperglycemia and high-fat diet (HFD)-induced hyperlipidemia (Figure 6A and Supplemental Material) in 134 (75 C57BL/6 and 59 ApoE^-/-^) mice. TTC staining at 24 h (Figure 6B) after UCAO with either saline or STZ showed that infarction did not occur in any C57BL/6 (0/6 and 0/8, respectively) or ApoE^-/-^ (1/10 and 0/11, respectively) mice. HFD without STZ administration did not cause infarction after UCAO in either C57BL/6 (0/12) or ApoE^-/-^ (0/9) mice. In contrast, STZ+HFD+UCAO induced large arterial infarction after UCAO in 22% (11/49, including three lethal infarction) of C57BL/6 mice and 31% (9/29) of ApoE^-/-^ mice. Approximately half the infarcted animals (6/11 C57BL/6 and 4/9 ApoE^-/-^ mice) had large infarcts. In the logistic regression analysis, fasting glucose levels (at 7 d before UCAO) and triglyceride levels (at 24 h after UCAO) were significant predictors of UCAO alone-mediated infarction occurrence by 24 h, after adjusting for the mouse strain (both *P*<0.005; Supplemental Table 1). Total cholesterol level was a marginally significant predictor (*P*=0.058). Subsequent experiments were performed to examine the involvement of NOS inhibition in UCAO-mediated infarction in hyperglycemic- hyperlipidemic mice (Figure 6C), blood levels of two endogenous NOSi (asymmetric dimethylarginine [ADMA] and SDMA) were measured in 20 (10 HFD+STZ and 10 vehicle control) C57BL/6 mice (without UCAO). SDMA levels (mean±SE) were significantly higher in the STZ+HFD group (26.8±1.5 ng/mL) than in the control group (17.1±1.2 ng/mL, *P*<0.001), which was not the case for ADMA (*P*=0.12).

**Figure 6.**
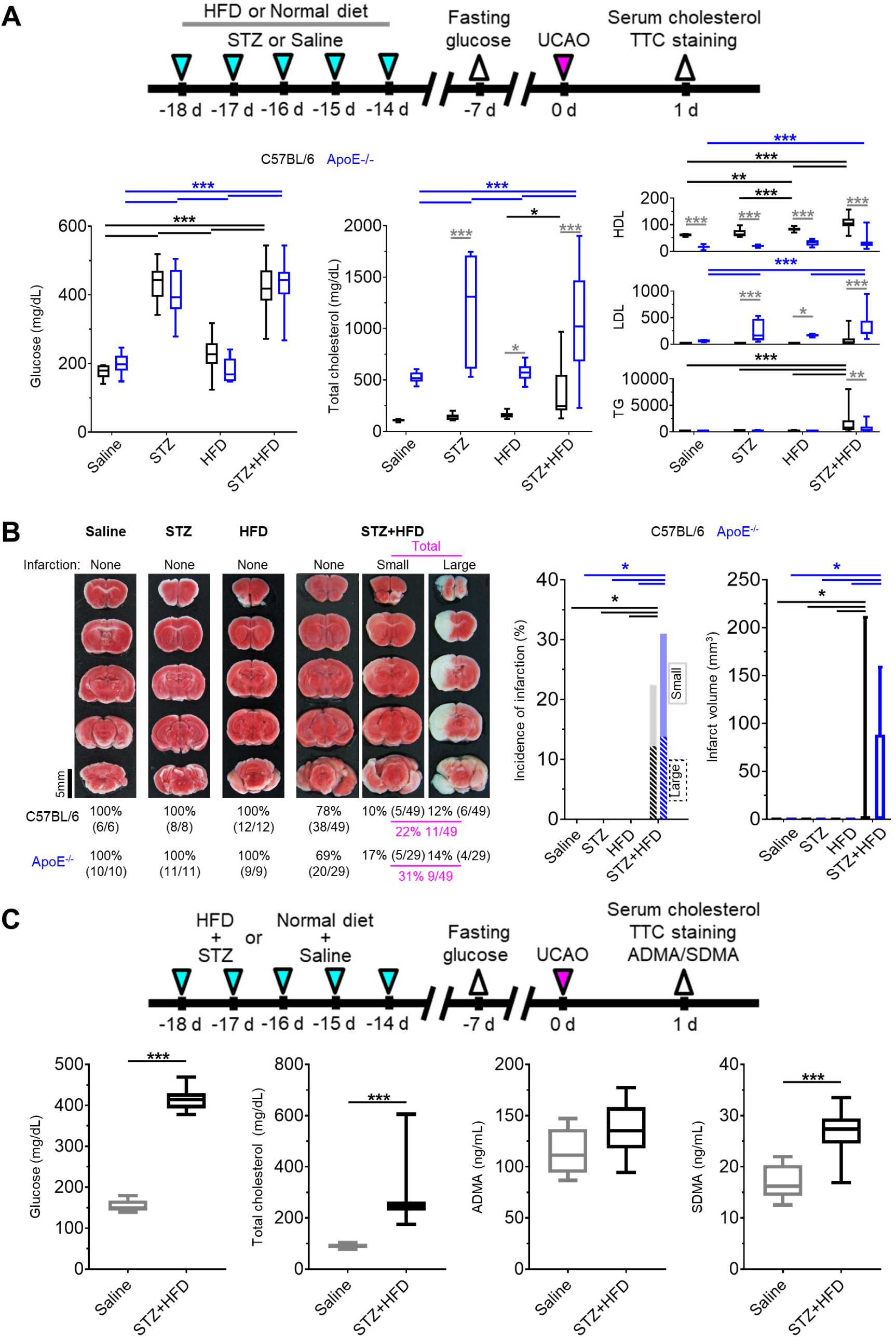
UCAO without L-NAME could induce acute large artery cerebral infarction in mice with hyperglycemia and hyperlipidemia. (**A**) Top, timeline for experiments on the effects of prior HFD and/or STZ treatment in C57BL/6 and ApoE^-/-^ mice receiving UCAO. Bottom, blood levels of glucose and cholesterol (total cholesterol, HDL cholesterol, LDL cholesterol, and TG) after HFD and/or STZ treatment: \**P*<0.05, \*\**P*<0.01, and \*\*\**P*<0.001 (black and blue: inter-group difference; gray: inter-strain difference), Two-way ANOVA and Sidak’s multiple comparisons for post-hoc tests. (**B**) Left, representative TTC staining images in each group, stratified by lesion volume (no, small [≤100 mm3], and large [>100mm3] infarction). Right, incidence of cerebral infarction and infarct volumes (box-and-whiskers plots). Infarctions occurred in the STZ+HFD groups only, regardless of strain. Lesion volumes did not differ significantly between the strains. (**C**) Top, timeline for experiments to examine the effects of HFD+STZ treatment on two endogenous NOSi (ADMA and SDMA) as well as of glucose and cholesterol. Bottom, measurement results (box-and-whiskers plots): \*\*\**P*<0.001, Mann-Whitney *U* test.

### Hyperglycemia and hyperlipidemia associated with infarct volume in patients with UCAO- mediated acute stroke

To obtain proof of principle that NOS dysfunction due to hyperglycemia and hyperlipidemia may be linked to infarction after carotid occlusions in humans, we conducted a multi-center clinical study using diffusion-weighted-MRI (DW-MRI) to identify factors that associate with the volume of acute (<7 d) cerebral infarction due to UCAO. We consecutively enrolled 438 patients (mean±SD age, 72±11 years), 271 (62%) of whom were male (Supplemental Table 2). Multiple linear regression analysis (Model 1 in Table 1) showed that fasting glucose and total cholesterol levels as well as initial stroke severity (admission National Institutes of Health Stroke Scale score) and atrial fibrillation were significantly associated with infarct volume (all *P*<0.05). In addition, age tended to be associated with infarct volume (*P*=0.07). After further adjusting for stroke subtype (Model 2 in Table 1), fasting glucose and initial stroke severity were significantly associated with infarct volume (both *P*<0.05), while total cholesterol and age tended to be associated with infarct volume (*P*=0.06 and 0.05, respectively).

**Table 1.**
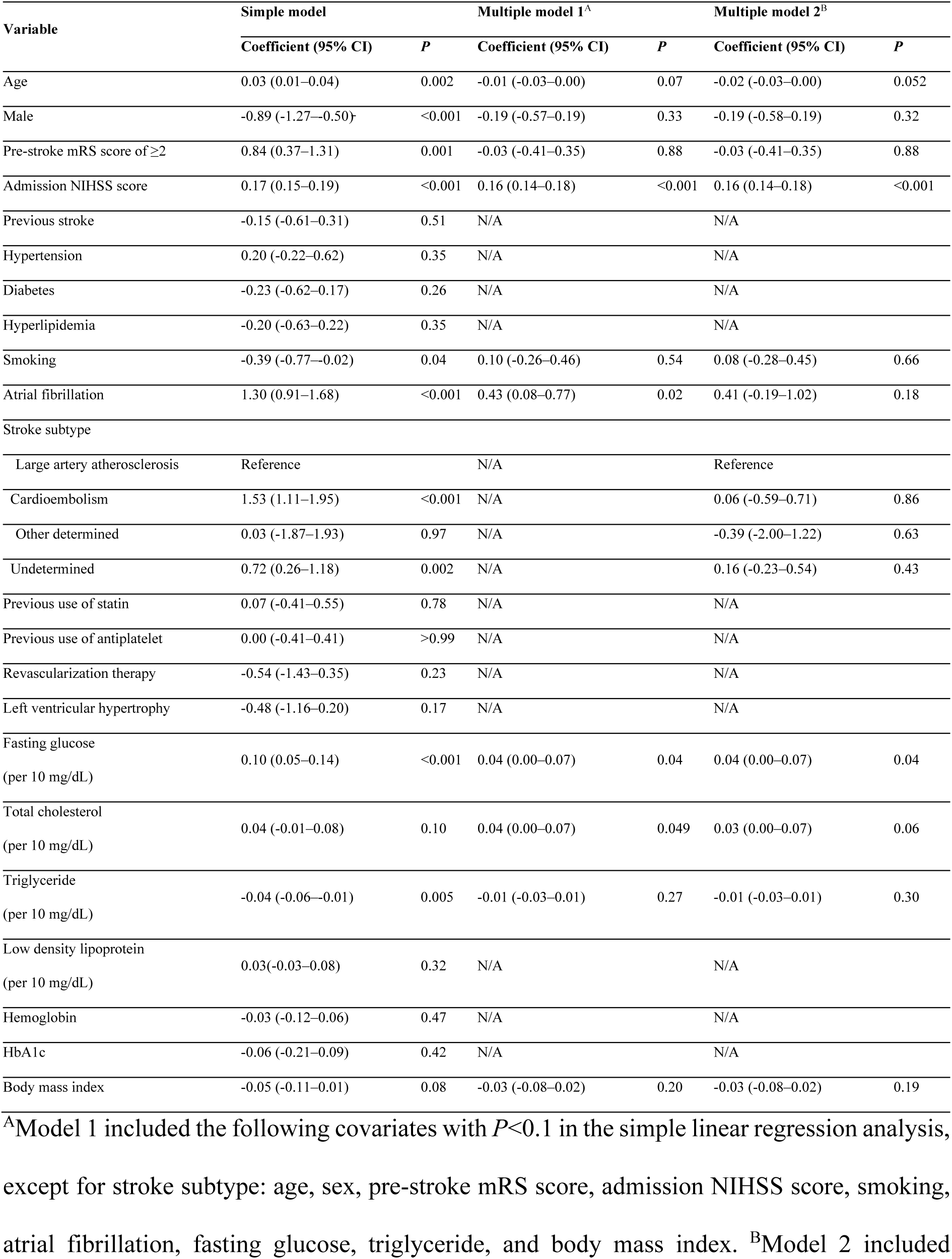
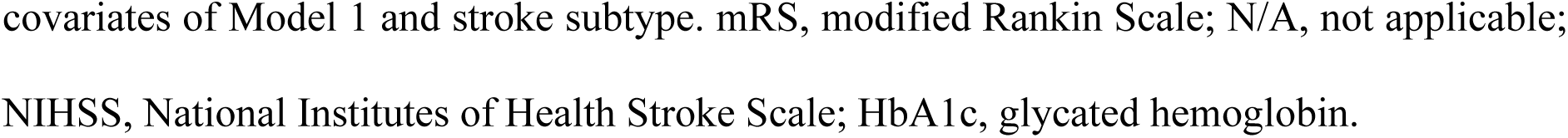
Linear regression analysis showing factors associated with log-transformed infarct volume in 438 patients with acute ischemic stroke due to occlusion of a proximal extracranial (internal or common) carotid artery.

### Mendelian randomization analysis identified a causal relationship between NOS inhibition and human ischemic stroke

To further support our hypothesis that NOS deficiency converts carotid occlusions into infarction, we performed Mendelian randomization analysis: an analytical approach that simulates a randomized controlled trial by using genetic variants (SNPs) as instrumental variables to control for unmeasured confounding and reverse causation (25). Specifically, we utilized i) GWAS summary statistics to select significant SNPs (Figure 7A) for three exposures (the main substrate for NOS, L-arginine, and the two endogenous NOSi, ADMA and SDMA) and for the outcome (ischemic stroke) and ii) previous large case-control studies of ischemic stroke with extensive genotyping (GIGASTROKE). We found SDMA concentration had a significant causal effect on risk of ischemic stroke using inverse variance weighted (IVW) method (Figure 7B): OR=1.24 (95% CI, 1.11-1.38), *P*=7.69×10^-5^. In contrast, no significant causal relationship could be inferred for either L-arginine or ADMA using IVW method. These findings were reproduced in sensitivity analyses (using different Mendelian randomization methods: Mendelian randomization-Egger [MR-Egger], weighted median, and weighted mode) that were conducted to account for genetic pleiotropy (Figure 7C). The sensitivity analysis could not be performed on L-arginine due to the lack of independent SNPs (n=2) qualified as instrumental variables (26).

**Figure 7.**
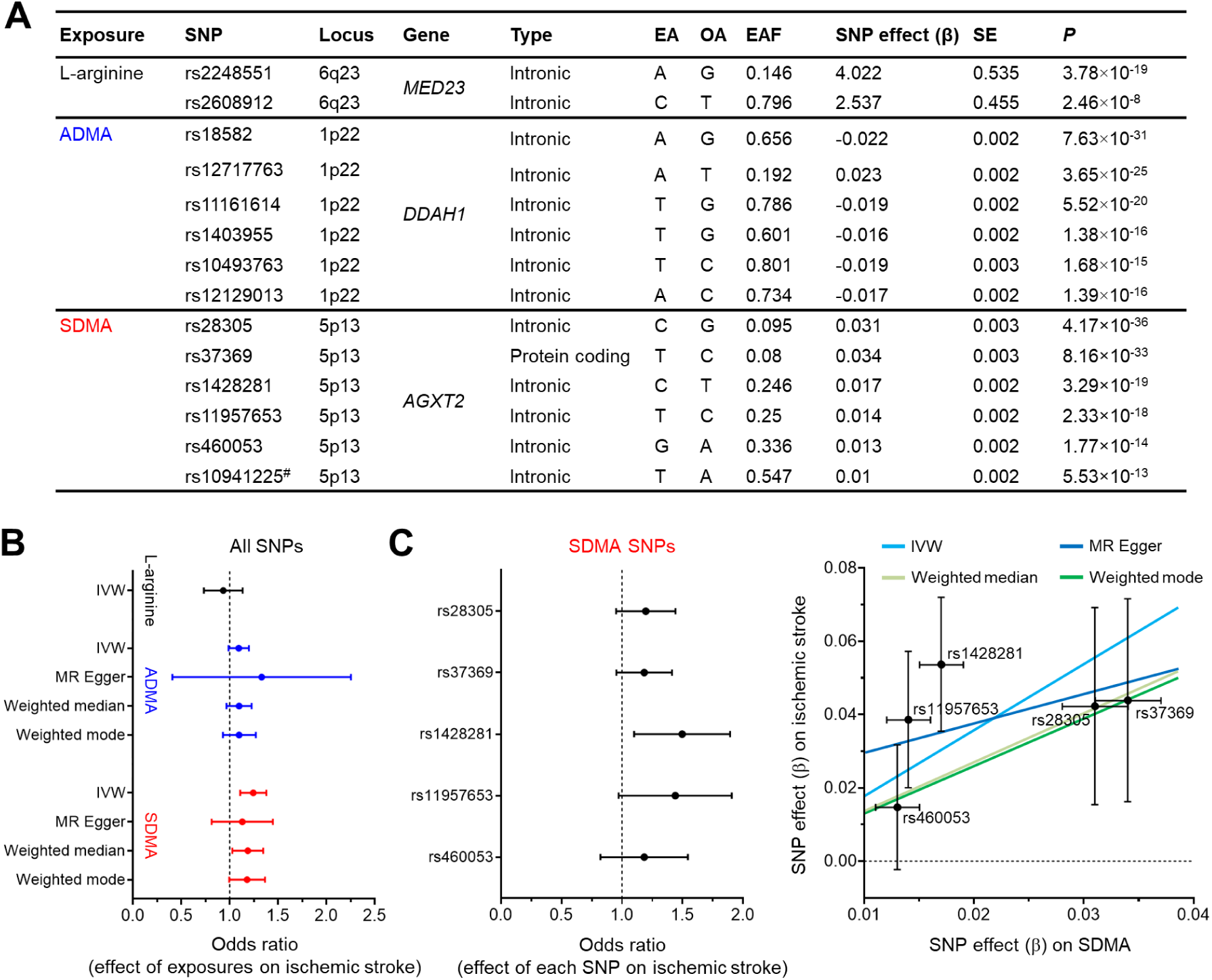
Mendelian randomization analysis showed a causative role of SDMA, an endogenous NOSi, in human ischemic stroke. (**A**) Characteristics of SNPs used in the Mendelian randomization study. # indicates an excluded SNP as a cross-correlated SNP, determined by linkage disequilibrium clumping (R^2^<0.01). EA, OA, and EAF indicate effect allele, other alleles, and effect allele frequency, respectively. SNP effect was calculated as increase in exposure (μmol/L) per EA. (**B**) Forest plot for the effect of exposures (L-arginine, ADMA, and SDMA) on risk of ischemic stroke, assessed by using different Mendelian randomization methods. OR was estimated using all SNPs for each exposure (per 1 SD increase in L-arginine [25.2 μmol/L], ADMA [0.13 μmol/L], and SDMA [0.12 μmol/L]). The error bars represent the 95% CIs. (**C**) Left, Forest plot for the effect of each SDMA SNP on risk of ischemic stroke. The error bars represent the 95% CIs. Right, Scatter plot for the effect of SDMA on risk of ischemic stroke. The error bars represent SE. Regression lines for four Mendelian randomization methods are shown.

## Discussion

This study demonstrates that UCAO-mediated benign oligemia can transform into hemispheric infarction when mice are treated either before or after UCAO with a single-dose intraperitoneal injection of a NOSi (L-NAME). This transformation was associated with SDI, which causes vasoconstriction by inverse neurovascular coupling under pathological conditions (19). These findings emphasize the importance of NO deficiency in rendering oligemic tissue vulnerable to development of SDI and cerebral infarction, as confirmed by successful rescue with the NO donor molsidomine. We also found that in the presence of hyperglycemia and hyperlipidemia, probably, in part, via increasing the endogenous NOSi SDMA, UCAO alone, without L-NAME administration, induced infarction in C57BL/6 mice and ApoE^-/-^ mice far more frequently (22.4% and 31.0%, respectively) than in naïve C57BL/6 mice (∼2%). In line with this animal research, subsequent clinical investigation showed that patients with hyperglycemia or hyperlipidemia had significantly larger infarcts in acute carotid occlusion stroke. Lastly, Mendelian randomization analysis revealed that SDMA plays a causative role in the risk of human ischemic stroke. Future research should advance the knowledge on the role of NOS inhibition and/or NO deficiency in the risk and severity of ischemic cerebrovascular disease and the underlying mechanisms. Such work could employ our new animal model used in the present study for *in vivo* imaging and drug testing.

Occluding MCA to reduce CBF to less than ∼20% of baseline is known to cause acute cerebral infarction in mice (15). A previous study using C57BL/6 mice (27) showed that i) spontaneous SDI occurred at 1–10 min following distal MCA occlusion (MCAO)-mediated CBF reduction to ∼45% of baseline and ii) recurrent SDIs could occur subsequently. The authors proposed that i) ischemia-induced SDI in the core triggers infarct formation and ii) subsequent SDIs in the oligemic penumbra (outside the core) expand the infarct. Our study showed that i) L-NAME+UCAO initially decreased CBF to an oligemic level (∼60%) in C57BL/6 and BALB/c mice with infarction (as assessed at 24 h), ii) their SDIs began in a more delayed fashion at ∼30–330 min (or possibly during the 6-24 h period when LSCI monitoring was not performed), and iii) pre-SDI rCoBF (at 3 min before the first SDI) was ∼45%. Thus, decreasing CoBF to ∼60% combined with NOSi-mediated reduction of NO might be, eventually, equivalent to ∼45% CoBF in terms of risk of acute cerebral infarction, thereby indicating NOSi plays a critical role in transforming oligemia to infarction in association with SDI. It is tempting to hypothesize that either avoiding risk factors that reduce NO availability or adopting clinical procedures or therapies that preserve or augment NOS activity could lessen or prevent the development of SDI and infarction of normal or oligemic brain tissue and ischemic penumbra.

SDI incidence was high in our study group. LSCI during the initial 6 h after L-NAME+UCAO revealed that SDI occurred in 11 of 27 mice that were later found either to have undergone cerebral infarction, as revealed when examined at 24 h, or to have died before that time-point due to large (lethal) infarction, particularly in BALB/c mice with relatively low pre- SDI rCoBF (Figure 2D and Supplemental Figure 3B). However, neither SDI nor death occurred in 17 non-infarcted animals. These results suggest that NOSi-mediated NO deficiency renders UCAO-mediated oligemic brain tissue vulnerable to SDI and that its propensity is higher when CoBF is lower within the oligemic CoBF range. As speculated above, during the 6 to 24 h period, SDI likely occurred in the other 16 of the 27 infarcted mice but not in the 17 non-infarcted mice. Further investigation is needed to determine if SDI is necessary for L-NAME+UCAO to generate cerebral infarction.

After UCAO, the timing of any disposition in NO availability appears to profoundly impact adaption to oligemia vis-à-vis evolution to infarction, since prolonged UCAO protected animals from acute ischemic stroke after L-NAME administration. Several factors could account for this protection. For example, inhibiting smooth muscle contraction/tone, if present in the posterior and anterior communicating collateral arteries, would increase CBF to the territory at risk. This could also result from vasodilation of secondary pial/leptomeningeal collaterals, which is known to occur within seconds-to-minutes after arterial obstruction due to fluid shear stress-mediated rise in eNOS activity (28). Also, primary (29) and pial (6) collaterals undergo expanded anatomic diameter (collateral remodeling) that, in the latter, begins within hours following obstruction and can double diameter by 36 h post-MCAO. These structural changes could lessen the L-NAME-mediated NOS inhibitory effect on CBF and thus infarction severity, as we observed, by decreasing the likelihood that SDI persists enough to cross the so-called commitment point. Alternatively or additionally, delaying L-NAME administration after UCAO might have allowed time for the development of increased neuronal resistance to ischemia or elevated neuronal tolerance to L-NAME or NO deficiency.

Arterial vasospasm, SDI, and microcirculatory dysfunction, which is linked to altered autoregulation and microvascular thrombosis, have all been implicated in cerebral infarction occurrence after aneurysmal subarachnoid hemorrhage (30). L-NAME+UCAO-mediated cerebral infarction was accompanied by all of these neurovascular alterations, as shown by LSCI and intravital microscopy, which revealed the involvement of SDI, arteriolar vasoconstriction, and microcirculatory dysfunction, such as altered autoregulation (i.e., relatively lower [∼40-50% rather than the usual ∼60-70%] post-UCAO rCoBF) and microvascular thrombosis (i.e., platelet – leukocyte stalls with sluggish capillary flow). Further, we demonstrated that the widely-used clinical drug cilostazol, which has both vasodilatory and anti-thrombotic effects (21, 22), could prevent or attenuate L-NAME+UCAO-mediated cerebral infarction, further supporting the pathogenic roles of both hemodynamic and thrombotic factors in this preclinical model. Note also that in the experiments using a heparinized catheter system to monitor heart rate and BP, infarction occurred relatively infrequently, 17% (2/12) and 36% (5/14) in C57BL/6 and BALB/c mice, respectively, possibly due to the anticoagulant locking solution leaking or spilling into the systemic circulation. Note that when only LSCI was performed without such cardiovascular monitoring (with heparin locking), infarction occurred more often in both mouse strains: 53% (10/19) and 64% (16/25), respectively.

In patients with acute carotid occlusion stroke, higher levels of glucose and cholesterol were significantly associated with larger infarctions. Moreover, in mice with hyperglycemia and hyperlipidemia, which was associated with ∼60% higher SDMA, UCAO alone, without L-NAME administration, induced large arterial infarction more than 10-times as often when compared to naïve mice. These clinical and preclinical findings of the present study align with previous basic and clinical research, showing that hyperglycemia (31) and hyperlipidemia (32, 33) can reduce eNOS activity and NO production. In non-diabetic patients with acute myocardial infarction (34), admission hyperglycemia was an independent predictor for long-term prognosis. Cholesterol levels, even within the normal range, may be inversely related to endothelial (NO-related) vasodilation (35). Future clinical research should confirm our preclinical findings, i.e., if i) NOS dysfunction both initiates acute ischemic stroke and aggravates pre-existing acute infarction in the setting of acute proximal artery occlusion and ii) NO donors prevent these outcomes.

Our Mendelian randomization analysis indicated SDMA-related inhibition of NOS activity (indirectly via limiting cellular uptake of the key substrate L-arginine) (36) plays a causal role in human ischemic stroke (OR=1.24). With regard to L-arginine and ADMA (an endogenous compound that competes with L-arginine for binding to NOS’s active site) (37), no significant link to ischemic stroke was found. These results accord with a previous Mendelian randomization study that linked higher SDMA (OR=1.07), but not the substrate L-arginine or the indirect NOSi ADMA, to elevated risk of ischemic heart disease. It has been suggested that supplementing L-arginine does not necessarily increase NO (38) or may raise the methylation demand for L-arginine, thereby counteracting antioxidant and antiapoptotic effects (39) by increasing NO (40).

To the best of our knowledge, prior to our study no molecule or drug that induces large cerebral infarction in the setting of mildly reduced CBF had been identified. Given the known protective role of eNOS in cerebral ischemia (41), we based our research on the *a priori* hypothesis that giving L-NAME intraperitoneally before or after inducing oligemia or mild cerebral ischemia would cause *de novo* cerebral infarction. Traditional MCAO models typically involve either inserting an intraluminal suture via the carotid artery to occlude the proximal portions of the MCA or performing craniotomies to expose and ligate or permanently electrocoagulate the distal MCA (42, 43). These models are well-established but require technically more demanding and time-consuming microvascular procedures, particularly in mice. Hence, there is a need for simpler, faster models.

Hattori et al. developed a new ischemic stroke mouse model that uses ameroid constrictors to gradually narrow and finally occlude both carotid arteries (44). At one month after the bilateral UCAO, H&E staining revealed cerebral infarcts. This model nicely reflects several characteristics of natural carotid artery occlusive stroke in humans; however, bilateral CAO is a rare vascular disease. Moreover, cerebral infarction did not develop until 7 d after the chronic UCAO procedure, which relied on costly constrictors; accordingly, this model may not be suitable for high-throughput animal research. Additionally, while large territorial infarction is typical of acute carotid stroke in humans (3), this model only produced small infarcts, and the detailed mechanism of their formation remains unclear. In contrast, the present study introduces a new mouse stroke model that may have several potential advantages.

Our model, which we term NOSi-mediated large Artery Ischemic stroke Model (NAIM), is technically less demanding, and it yields ∼40 mice per day per surgeon, whereas the widely employed intraluminal suture model usually yields ∼5 mice. Single-dose molsidomine pre-treatment or post-treatment lessened cerebral infarction and stroke-related death (i.e., lethal infarction) in NAIM. Thus, our model provides a preclinical *in vivo* assay for screening molsidomine-like molecules or drugs that could either counteract or exacerbate NO deficiency- or SDI-related pathogenesis in stroke or other diseases involving reduced NO bioavailability. In addition, our findings support the concept that molsidomine might be useful in preventing or treating ischemic stroke caused by large artery steno-occlusion, thereby warranting further investigation to confirm this hypothesis. Furthermore, employing NAIM allowed for vasodilation-related therapeutic distinctions among broadly used antiplatelet drugs (cilostazol vs. aspirin and clopidogrel), as previously demonstrated (22) by utilizing a photothrombotic MCAO model, which is superior to mechanical MCAO models for assessing thrombotic stroke.

In addition to presenting our new model, our study has multiple strengths. First, the model is well characterized, including its mechanism, i.e., the contribution of NOSi-mediated NO deficiency to SDI and infarction. Second, the procedures can be performed quickly and the technique is surgically straight-forward, so the NAIM model may allow for high-throughput studies. Third, most animals had either territorial large artery infarction or no infarction at all, so these dichotomized tissue outcomes will be statistically easy to analyze; the absence of infarction in this model does not mean technical failure. Fourth, the heterogeneity of NAIM-related outcomes even within the same mouse strain may provide opportunities to clarify genetic or molecular determinants of oligemia- or penumbra-to-infarct transition (45), compare the biology of fast vs. slow progressors with large vessel occlusion (46), and investigate novel pathophysiologic mechanisms underlying the well-known wide spectrum of stochastic differences in clinical stroke outcomes. Fifth, including the Mendelian randomization analysis provides a more rigorous and unbiased understanding of the causal relationship between NOS inhibition and the risk of cerebral infarction in humans, thereby supporting our preclinical model-based data and, in conjunction with our multi-center clinical data, emphasizing the clinical implications of the current study.

Nevertheless, there are a few caveats and further questions remain. First, NAIM has a relatively high mortality rate, which may lead to needing large numbers of animals when testing complex hypotheses. In this respect, the strength of NAIM as a high-throughput preclinical model of large artery ischemic stroke is worth of noting. Second, further investigation is required regarding what contributes, independently and interactively, to the mouse strain differences in response; these three commonly used strains may have varied vulnerability to excitotoxicity (47) in addition to having different collateral status (48). Third, our Mendelian randomization study could have been biased due to unexpected pleiotropic effects of genes, despite the consistent results of various sensitivity analyses.

In conclusion, we report novel findings that NOS inhibition can transform UCAO- mediated oligemia into *de novo* hemispheric infarction, via SDI and arteriolar constriction/microvascular thrombosis. Our preclinical discovery is supported by subsequent bench-to-clinic and clinic-to-bench investigations: i) multi-center MRI research of 438 patients with UCAO, ii) animal experiments using mice with hyperglycemia/hyperlipidemia, and iii) Mendelian randomization analysis. Collectively, these inquires indicate NOS inhibition plays a causal role in the occurrence and progression of animal and human ischemic stroke. In addition, our data may warrant clinical investigations to confirm whether SDMA predicts the progression of brain oligemia to infarction following acute proximal artery occlusion and whether molsidomine protects against it. The present study also shows that NAIM, a new animal model of stroke, enables investigators to generate large artery cerebral infarction at high throughput, unlike traditional MCAO models that require technically more demanding and time-consuming microvascular procedures.

## Methods

### Sex as a biological variable

In the preclinical study, main results were derived using only male mice to avoid the confounding effect of the hormone cycle in females. Only one experiment included female mice to compare their results with those from male mice. In the clinical study, all statistical analyses accounted for sex as a biological variable by adjusting for it as a covariate.

### Study design for preclinical research

A total of 900 (∼11–12-week-old) C57BL/6 (n=517), BALB/c (n=257), SV129 (n=50), and ApoE^-/-^ (n=76) mice (DBL Co, Incheon, Korea) were used. Mice were fed *ad libitum* in a specific pathogen-free and climate-controlled environment maintained at 23±2°C and 50±10% humidity (mean±SD), with 12 h light-dark cycle. In the following experiments, after animals were either euthanized at designated time-points or found dead, their brains were subjected to *ex vivo* green-channel autofluorescence imaging (that we previously developed) (49) and TTC staining (50) to measure the volume of infarction (if any) and obtain autopsy evidence of stroke-related death (lethal infarction). Anesthesia was performed using 2% isoflurane through an inhalation mask, and body temperature was monitored using a rectal probe and maintained at 36.5°C.

Experiment 1 was performed after completion of a dose-finding pilot study (data not shown) to determine the single intraperitoneal dose of L-NAME as 400 mg/kg. A midline neck incision was made and the perivascular tissue was dissected to prepare for right common carotid artery ligation in C57BL/6 (n=140) and BALB/c (n=102) mice. Seven C57BL/6 mice, 12 C57BL/6 mice, and 75 (n=47 C57BL/6 and 28 BALB/c) mice, respectively, served as sham surgery group, L-NAME only group, and UCAO only group (for 24 h in 37 C57BL/6 and 14 BALB/c mice vs. for 7 d in 10 C57BL/6 and 14 BALB/c mice). The remaining 148 (n=74 C57BL/6 and 74 BALB/c) mice received both L-NAME and UCAO (L-NAME+UCAO) as follows: After L-NAME was administered, right UCAO was performed. In 45 (n=19 C57BL/6 and 26 BALB/c) mice randomly selected from the 148 L-NAME+UCAO group animals, we also performed LSCI (Moor Instruments, Axminster, UK), as previously described (51), to monitor relative CoBF for 6 h after UCAO. These 45 mice were euthanized 24 h later. The other 103 mice were euthanized at 1, 2, 3, and 6 d after UCAO. One BALB/c mouse that underwent LSCI was excluded for LSCI analysis because UCAO reduced rCoBF in the core region (ROI-1) by ∼90% (to 9.1% of the pre-L-NAME+UCAO baseline), which alone can cause cerebral infarction. In addition to the aforementioned experiments, we examined L-NAME+UCAO in SV129 mice (n=50), a strain known to have good posterior communicating and pial collaterals (48), at 1, 2, 3, and 6 d after UCAO; two mice were excluded due to death from undetermined cause. We also studied 22 female C57BL/6 mice, which underwent L-NAME+UCAO and were euthanized 24 h later.

Experiment 2 examined the effect of administration of L-NAME after – rather than before – UCAO at 3 h and 1, 2, 3, 5, and 7 d post UCAO in 108 C57BL/6 and BALB/c mice (n=9 per time-point per strain). Mice were euthanized 24 h after L-NAME administration.

Experiment 3 used a rescue design wherein 200 mg/kg of the NO donor, molsidomine (Sigma, St. Louis, MO, USA), and saline (330 µl) were administered intraperitoneally 30 min after L-NAME+UCAO to 88 mice (n=22 C57BL/6 and 22 BALB/c per treatment), which were euthanized 24 h later.

Experiment 4 simultaneously monitored BP (using a femoral arterial catheter and Small Animal Physiological Monitoring System; Harvard Apparatus, Holliston, MA, USA) and rCoBF (with LSCI) before (for ∼10 min) and after (for ∼90 min) 200 µl saline administration only, L-NAME administration only, UCAO only, or L-NAME+UCAO in C57BL/6 (n=27) and BALB/c mice (n=29).

Experiment 5 investigated whether L-NAME+UCAO causes macrovascular/microvascular thrombosis or thromboembolic steno-occlusion in 47 mice by performing: i) high-resolution *in vivo* microCT-based direct thrombus imaging serially at 0, 1, 2, 3, 4, 5, and 6 d after L-NAME+UCAO (n=9 C57BL/6 and 9 BALB/c mice); ii) intravital two-photon microscopy imaging through a cranial window at baseline and serially by 4 or 24 h after either left UCAO only (n=1 BALB/c mouse) or L-NAME + left UCAO (n=5 C57BL6 and 18 BALB/c mice); and iii) H&E staining of brain tissue sections at 24 h after L-NAME+UCAO (n=5 C57BL/6 mice). For further details, see Supplemental Material.

Experiment 6 explored the protective effects of three widely prescribed antiplatelet medications (vs. 200 µl saline control) on L-NAME+UCAO-mediated infarction in 96 C57BL/6 mice (n=24/group) pre-treated at a dose shown to have antithrombotic effects in rodents (22): aspirin 100 mg/kg/day (Bayer, Leverkusen, Germany), clopidogrel 10 mg/kg/day (Plavix; Sanofi, Paris, France), or cilostazol 100 mg/kg/day (Pletaal; Korea Otsuka Pharmaceutical, Seoul, Korea) was orally administered once daily (twice a day for cilostazol) for 8 d, with the final dose given 1 h prior to L-NAME+UCAO. Mice were euthanized 6 d later. Experiment 7 queried if UCAO without L-NAME administration could induce infarction when mice (n=92 C57BL/6 and 76 ApoE^-/-^) had STZ-mediated hyperglycemia and/or HFD- mediated hyperlipidemia, considering the reported link of these cardiovascular risk factors with NOS dysfunction (23, 24). In addition to glucose and cholesterol levels, blood levels of two endogenous NOSi (ADMA and SDMA) were measured in 20 (10 HFD+STZ and 10 vehicle control) of the C57BL/6 mice (without UCAO) to examine the involvement of NOS inhibition with subsequent UCAO-mediated infarction. For further details, see Supplemental Material.

### Analysis of LSCI data

As shown in Supplemental Figure 1, we defined three ipsilesional ROIs (each diameter ∼1.3 mm) on the cerebral cortex: ROI-1 (oligemic core), ROI-2 (anteromedial to ROI-1), and ROI-3 (parietal association cortex; anteromedial to ROI-2), taking into account post-UCAO low rCoBF territories on LSCI. ROI-6, -5, and -4 were respectively defined on the corresponding positions of the contralateral hemisphere. After quantifying the 6 h of LSCI monitoring data by calculating the flux values in the six ROIs of each video frame (1-60 frames/min), we computed each rCoBF value as a percentage relative to the pre-L-NAME+UCAO baseline, thereby generating rCoBF values. The occurrence of SDI was confirmed by both visual inspection and quantification of the LSCI datasets. To clarify the link between SDI and subsequent infarction, we stratified all mice into the following four groups by SDI occurrences (in ROI-1 during the 6 h or 1.5 h LSCI monitoring) and cerebral infarction (up to 24 h). For further details, see Supplemental Material.

### Statistical analysis for preclinical research

Data are presented as mean±SE, median (IQR), and number (percentage), as appropriate. Small or large infarction were defined as lesion volume smaller or larger (52) than 100 mm^3^ on TTC staining. Lethal infarction was defined as a TTC-confirmed infarction that resulted in stroke-related premature death before predetermined sacrifice. Severe infarction was defined as a composite outcome of large and lethal infarction. We used chi-square test or Fisher’s exact test, as appropriate, for categorical variables. We used Student’s *t* test, Mann-Whitney *U* test, or Kruskal-Wallis test with post-hoc pairwise Dunn’s tests for continuous variables, as appropriate. We used survival curves, obtained by the Kaplan-Meier method, and log-rank test with post-hoc pairwise comparisons to compare cumulative death rates between groups. To assess inter-group differences in rCoBF values (fixed effects) by comparing the repeated measures among time-points or groups while accounting for variability due to random effects (subject-specific intercepts), we obtained LS means, which are estimated marginal means from linear mixed models that accounts for the correlation structure inherent in repetitively measured data (53), and performed pairwise post-hoc tests with Sidak’s multiple comparison adjustment. We performed logistic regression analysis to identify predictors of infarct occurrence after UCAO in STZ-treated and/or HFD- fed mice. Data were analyzed using SPSS software 18.0 (IBM SPSS, Chicago, IL, USA) and SAS 9.4 (SAS Institute Inc, Cary, NC, USA). All graphs were constructed by using GraphPad Prism 10 (GRAPH PAD software Inc, San Diego, CA, USA).

### Study design for clinical research

The objective of the following two clinical studies were to corroborate animal study findings.

To identify risk factors that associate with the volume of acute cerebral infarction due to proximal CAO, we utilized the Korean image-based stroke database (54–57) of a prospective multi-center project, in which 11 stroke centers in Korea participated from May 2011 to February 2014. The institutional review boards of all participating centers approved this project, and patients or their legally authorized representatives provided written informed consent. A total of 438 consecutive patients with acute ischemic stroke due to occlusion of the extracranial proximal (internal or common) carotid artery were included in the final analysis, after exclusion of the following 226 patients who i) received revascularization (intravenous thrombolysis or endovascular thrombectomy) therapy before MRI (n=207), ii) did not undergo MRI (n=11), iii) had poor-quality or unavailable DW-MR images (n=4), or iv) had MRI segmentation/registration-related errors (n=4). DW-MRI was performed on 1.5T (n=378) or 3.0T (n=39) MRI systems. Infarct volumes were quantified and converted to percentage lesion volumes (percentages of the total brain parenchymal volume), followed by log-transformation, as we previously published (55, 57). Data are presented as mean±SD, median (IQR), and number (percentage), as appropriate. Missing data for fasting glucose (n=26), glycated hemoglobin (n=84), total cholesterol (n=11), height (n=36), and weight (n=10) were replaced with the median value of the entire population. We used multiple linear regression models (with or without adjustment for stroke subtype) to predict infarct volumes using covariates with *P*<0.1 in the simple linear regression analyses. A two-sided *P*<0.05 was considered significant. Data were analyzed using the STATA 18.0 (Stata Corp, College Station, USA).

To estimate the causal effect of a NOS substrate and the two endogenous (direct and indirect) NOSi on risk of ischemic stroke, Mendelian randomization analysis was performed by using the “TwoSampleMR” package in R software (Version 4.3.2). Based on a previous Mendelian randomization study of ischemic heart disease (58), genetic instruments for the three exposures (the substrate L-arginine and the inhibitors ADMA and SDMA) and for the outcome (ischemic stroke) were derived from large-scale GWASs by selecting SNPs that met the genome-wide significance level (*P*<5×10^−8^). To ensure the two-sample nature of the analysis, the exposure datasets and the outcome datasets were taken from different GWASs. To exclude cross-correlated SNPs, linkage disequilibrium clumping was performed with the threshold of R^2^<0.01. Thus, 2, 6, and 5 SNPs were selected as instrumental variables for L-arginine, ADMA, and SDMA, respectively. Then, the primary analysis was completed using the IVW method. In addition, multiple sensitivity analyses were conducted with the Mendelian randomization-Egger (MR-Egger) (59), weighted median (60), and weighted mode (61) methods to account for genetic pleiotropy and to enhance the robustness of the inferred causal relationships.

### Study approval

Animal experiments were approved by the Institutional Animal Care and Use Committee at Dongguk University Ilsan Hospital. The Korean image-based stroke database (54–57) collected brain imaging and clinical data under the approval of the institutional review boards of all participating centers (DUIH2010-01-083-020), and patients or their legally authorized representatives provided written informed consent.

## Data availability

The data that support the findings of this study are available from the corresponding author upon reasonable request.

## Author contributions

DEK, DS, SJL, DSG, JS, EHL, JF, and CA conceived, designed, and planned the study. HK, JC, JWK, DS, SJL, HJJ, IP, TK, DSG, JSL, SHH, and KHJ acquired the data. JC, DEK, SJL, DSG, JSL contributed to quantitative and statistical data analyses. All authors contributed to interpretation of data, writing of the manuscript and critical review and revision of the manuscript. All authors had full access to all of the data in the study. DEK had final responsibility for the decision to submit for publication.

## Acknowledgements

This study was supported by the National Priority Research Center Program Grant (NRF-2021R1A6A1A03038865, to DEK) and the Basic Science Research Program Grant (NRF-2020R1A2C3008295, to DEK) of National Research Foundation, funded by the Korean government. The authors appreciate DHH and all members of the Clinical Research Collaboration for Stroke Korea-5 for their contributions to this study.

