## Supplemental Material for "Inhibition of nitric oxide synthase transforms carotid occlusion-mediated benign oligemia into *de novo* large cerebral infarction"

Supplemental Tables 1–2

Supplemental Figures 1–9

Supplemental Videos 1–6

Supplemental Results

Supplemental Methods

Supplemental References 1–9

**Supplemental Table 1. Logistic regression analysis showing predictors of infarct occurrence in mice receiving UCAO without L-NAME administration**

| Variable | OR (95% CI) | *P* |
|---|---|---|
| Strain | 0.14 (0.02–0.85) | 0.032 |
| Fasting glucose (per 10 mg/dL) | 1.01 (1.03–1.18) | 0.003 |
| Total cholesterol (per 10 mg/dL) | 0.98 (0.96–1.00) | 0.058 |
| Triglyceride (per 10 mg/dL) | 1.01 (1.00–1.01) | 0.002 |

**Supplemental Table 2. Baseline characteristics of 438 patients with acute ischemic stroke due to occlusion of a proximal extracranial (internal or common) carotid artery**

| Variable | Stroke patients (n=438) |
|---|---|
| Mean age (SD), years | 71.7 (11.2) |
| Male, n (%) | 271 (61.9) |
| Pre-stroke mRS score of ≥2, n (%) | 86 (19.6) |
| Mean admission NIHSS score (SD) | 10.1 (7.6) |
| Previous stroke, n (%) | 99 (22.6) |
| Hypertension, n (%) | 312 (71.2) |
| Diabetes, n (%) | 156 (35.6) |
| Hyperlipidemia, n (%) | 118 (26.9) |
| Smoking, n (%) | 208 (47.5) |
| Atrial fibrillation, n (%) | 149 (34.0) |
| Stroke subtype, n (%) | |
| Large artery atherosclerosis | 192 (43.8) |
| Cardioembolism | 140 (32.0) |
| Other determined | 4 (0.9) |
| Undetermined | 102 (23.3) |
| Previous use of statin, n (%) | 85 (19.4) |
| Previous use of antiplatelet, n (%) | 135 (30.8) |
| Revascularization therapy, n (%) | 21 (4.8) |
| Left ventricular hypertrophy, n (%) | 37 (8.5) |
| Mean fasting glucose (SD), mg/dL | 125.3 (41.0) |
| Mean total cholesterol (SD), mg/dL | 172.5 (42.5) |
| Mean triglyceride (SD), mg/dL | 114.0 (77.4) |
| Mean low density lipoprotein (SD), mg/dL | 109.2 (37.1) |
| Mean hemoglobin (SD), g/dL | 13.5 (2.1) |
| Mean HbA1c (SD), % | 6.5 (1.4) |
| Mean height (SD), cm | 162.2 (8.4) |
| Mean weight (SD), kg | 60.6 (11.0) |
| Mean body mass index (SD), kg/m$^2$ | 22.9 (3.3) |
| Median Infarct volume (IQR), % of brain | 1.0 (0.3-5.8) |

Data are mean (SD), number (percentage), or median (IQR). Some data were missing for fasting glucose (n=26), HbA1c (n=84), total cholesterol (n=11), triglyceride (n=12), low density lipoprotein (n=37), height (n=36), and weight (n=10); these were replaced with the median of the entire population. mRS, modified Rankin Scale; NIHSS, National Institutes of Health Stroke Scale; HbA1c, glycated hemoglobin.

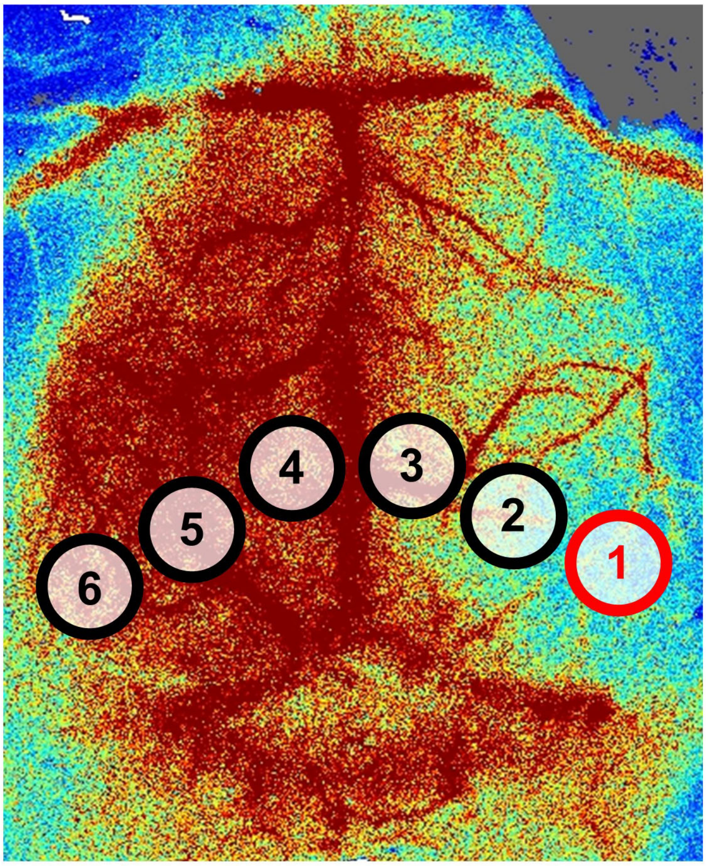


**Supplemental Figure 1. ROIs for quantitative analysis of LSCI data.** Six ROIs (diameter ~1.3mm) were placed on a representative LSCI to cover each hemisphere: from the anteromedial brain regions (ROI-3 in the right hemisphere vs. ROI-4 in the left hemisphere) of sensorimotor (primary somatosensory cortex, primary motor cortex, and secondary motor cortex) cortices (about 0-3 mm posterior and 0-1.3 mm lateral from the Bregma) to the posterolateral brain regions (ROI-1 in the right hemisphere vs. ROI-6 in the left hemisphere) including the secondary visual cortex and temporal association cortex (about 0.8-5.2 mm posterior and 3.2-4.4 mm lateral from the Bregma). The red-colored ROI-1 represent an oligemic core region in this C57BL/6 mouse with right UCAO after a single intraperitoneal dose of L-NAME. An experienced research assistant (DHH), who was blinded to experimental groups, made a small adjustment to ROI size and placement in each mouse, based on the size and shape of its brain.

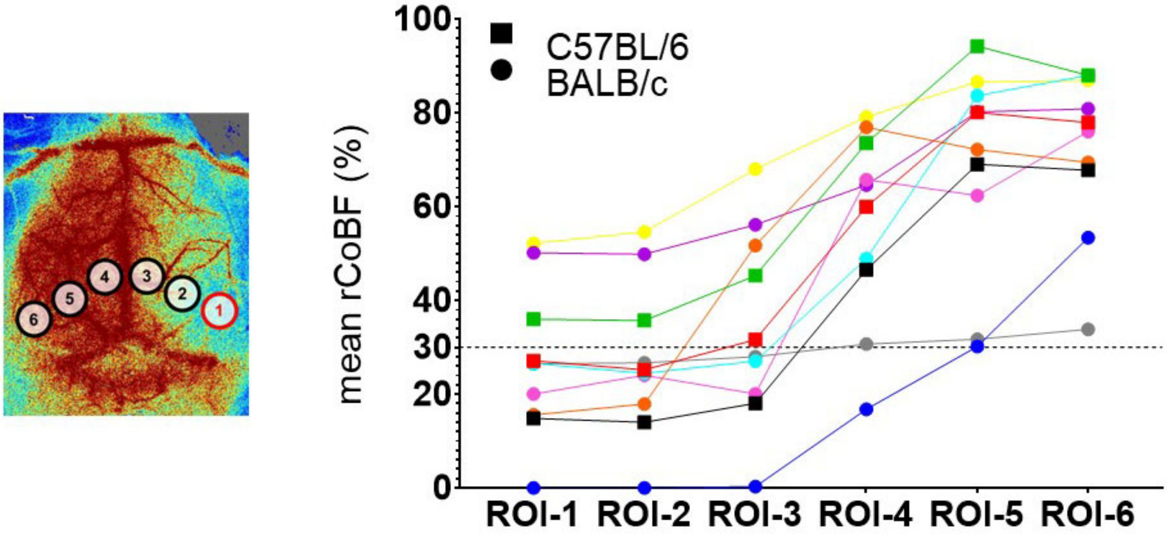


**Supplemental Figure 2. rCoBF (mean of the lowest values from individual SDI events) in every ROI of each SDI(+) mouse.** rCoBFs are lower in the ipsilateral hemisphere (ROI-1, 2, and 3) than in the contralateral hemisphere (ROI-4, 5, and 6).

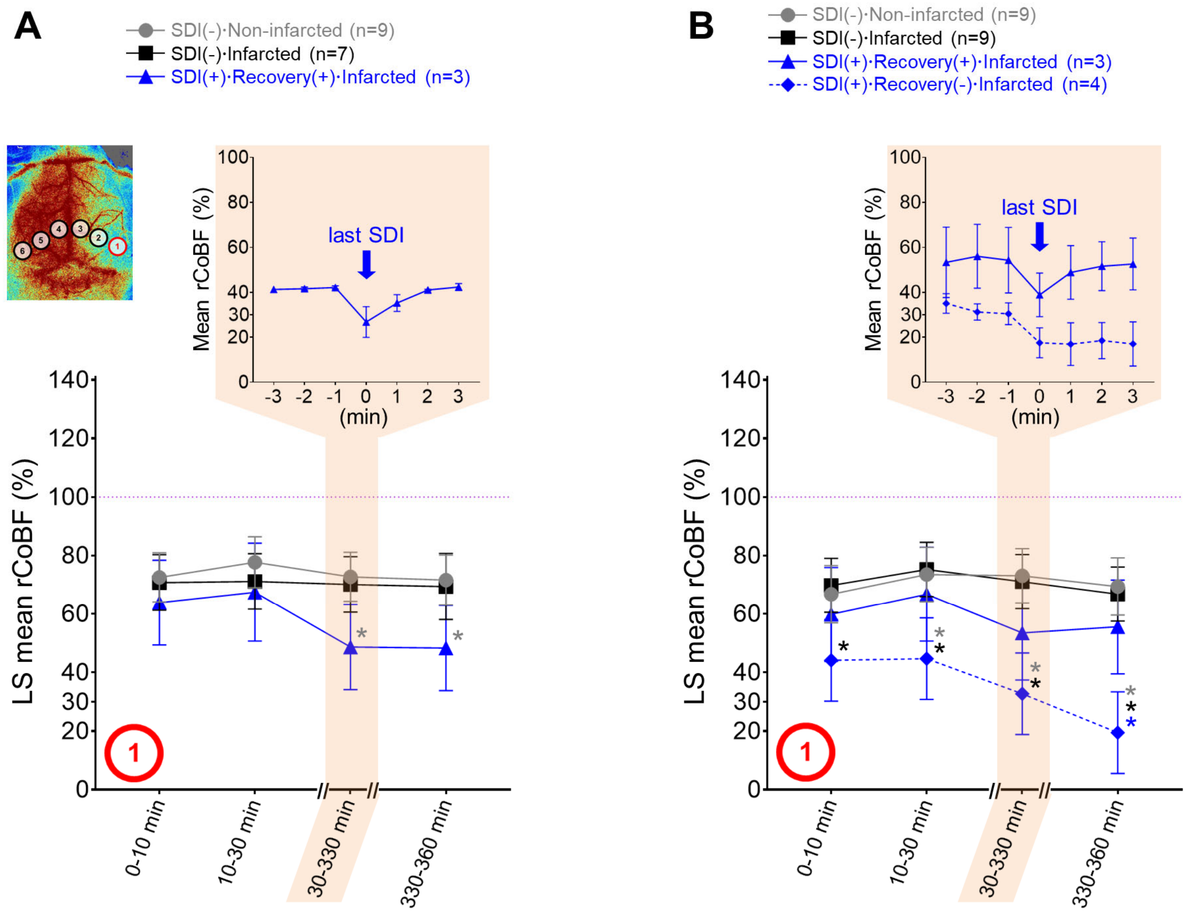


**Supplemental Figure 3. L-NAME+UCAO-mediated changes in the core rCoBF of C57BL/6 mice vs. BALB/c mice, stratified by the occurrence of SDI (up to 6 h) and cerebral infarction (up to 24 h).** (**A** and **B**) rCoBF, measured by LSCI after UCAO in (**A**) 19 C57BL/6 mice and (**B**) 25 BALB/c mice, pre-treated with a single intraperitoneal dose of L-NAME. We pre-specified four time periods (0-10, 10-30, 30-330, and 330-360 min) for the 6 h LSCI monitoring and calculated their mean rCoBF values (% of baseline, see Methods in the main text) in the core region (ROI-1). Pairwise comparisons with pre-specified multiple comparison correction were carried out using linear mixed models with random intercepts; LS mean values with 95% CIs are presented in the graphs. Gray, black, and blue * indicate *P*<0.05 vs. the SDI(-)·Non-infarcted group, SDI(-)·Infarcted group, and SDI(+)·Recovery(+)·Infarcted group, respectively, at each time period. Note that baseline (group mean±SE) values were not included in the statistical analyses of repeated continuous outcome measures. In addition to the statistical analyses, rCoBF values for the last SDI during the 30-330 min period are also displayed for each SDI(+) group (group mean±SE) in order to show SDI-related rCoBF drop to a trough level (inset graph in the shaded area).

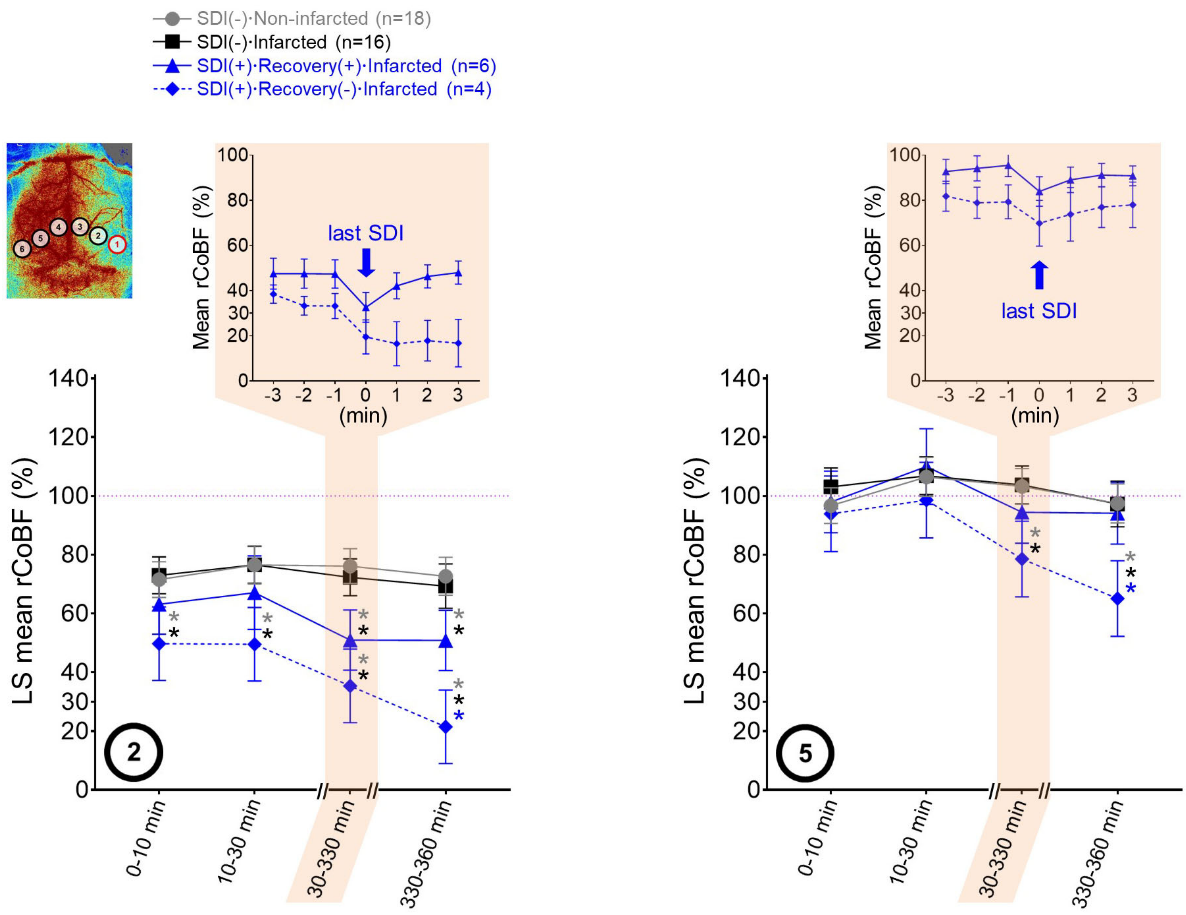


**Supplemental Figure 4. L-NAME+UCAO-mediated changes in the rCoBF of ROI-2 and ROI-5 of 44 mice, stratified by the occurrence of SDI (up to 6 h) and cerebral infarction (by 24 h).** rCoBF measured by LSCI after UCAO in 19 C57BL/6 mice and 25 BALB/c mice, pre-treated with a single intraperitoneal dose of L-NAME. We pre-specified four time periods (0-10, 10-30, 30-330, and 330-360 min) for the 6 h LSCI monitoring and calculated their mean rCoBF values (% of baseline, see Methods in the main text) in the ROI-2 and ROI-5. Pairwise comparisons with pre-specified multiple comparison correction were carried out using linear mixed models with random intercepts. Gray, black, and blue * indicate $P<0.05$ vs. the SDI(-)·Non-infarcted group, SDI(-)·Infarcted group, and SDI(+)·Recovery(+)·Infarcted group, respectively, at each time period. Note that baseline (group mean±SE) values were not included in the statistical analyses of repeated continuous outcome measures. LS mean values with 95% CIs are presented in the graphs. In addition to the statistical analyses, rCoBF values for the last SDI during the 30-330 min period are also displayed for each SDI(+) group (group mean±SE) in order to show SDI-related rCoBF drop to a trough level (inset graph in the shaded area). ROI-2 data showed mixed ROI-1 and ROI-3 characteristics, while ROI-5 data showed mixed ROI-4 and ROI-6 characteristics (see Figure 2D and Results in the main text).

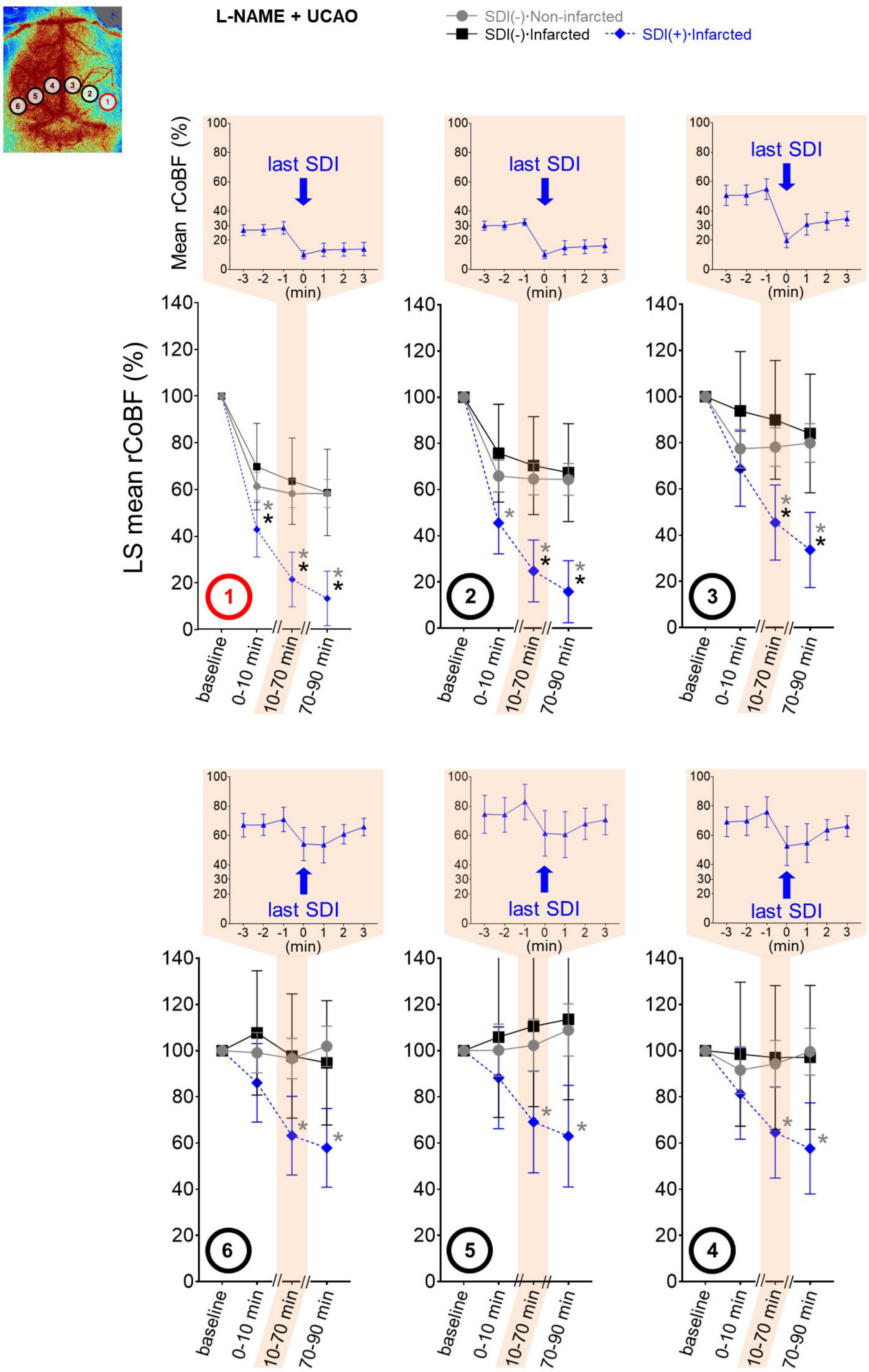


**Supplemental Figure 5. L-NAME+UCAO-related changes in all regional rCoBFs.** Heart rate, systolic and diastolic BP, and rCoBF (all as mean±SE) in all ROIs of 26 mice in L-

NAME+UCAO group. We stratified mice into four subgroups by the occurrence of SDI (with or without rCoBF recovery up to 90 min) and cerebral infarction (up to 24 h). Given that every SDI event observed during the 90 min monitoring period occurred between 10.2-70.0 min after L-NAME+UCAO, we pre-specified three time periods (0-10 min, 10-70 min, 70-90 min) and calculated their mean heart rate, BP, and rCoBF values for subgroups in each group. Pairwise comparisons with pre-specified multiple comparison correction were carried out using linear mixed models with random intercepts. Gray and black * indicates $P<0.05$ vs. the SDI(-)·Non-infarcted group and SDI(-)·Infarcted group, respectively, at each time period. LS mean values with 95% CIs are presented in the graphs. In addition to the statistical analyses, rCoBF values for the last SDI during the 10-70 min period are displayed for each SDI(+) group (as group mean±SE) in order to show SDI-related rCoBF drop to a trough level (inset graph in the shaded area). Corresponding heart rate and BP data (at the time-point of lowest rCoBF) are also presented in inset graphs.

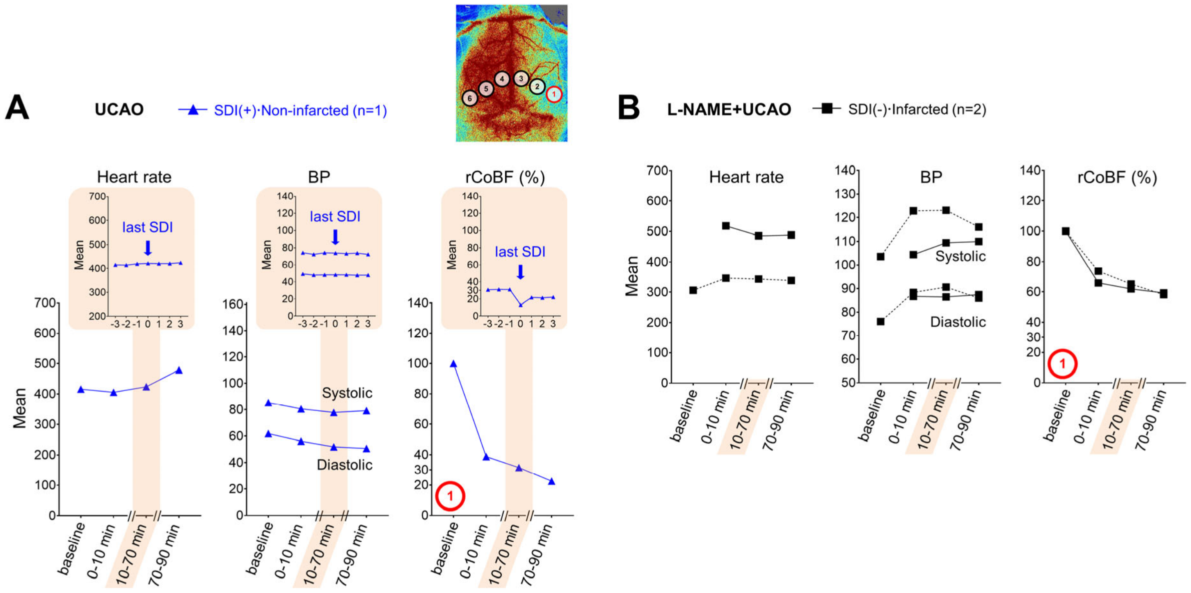


**Supplemental Figure 6. L-NAME+UCAO-related changes in heart rate, BP, and core rCoBF of minor group animals.** (**A** and **B**) Heart rate, systolic and diastolic BP, and rCoBF (all as mean±SE) in the oligemic core (ROI-1) of the three mice that are not presented in Figure 3: (**A**) UCAO only group and (**B**) L-NAME+UCAO group.

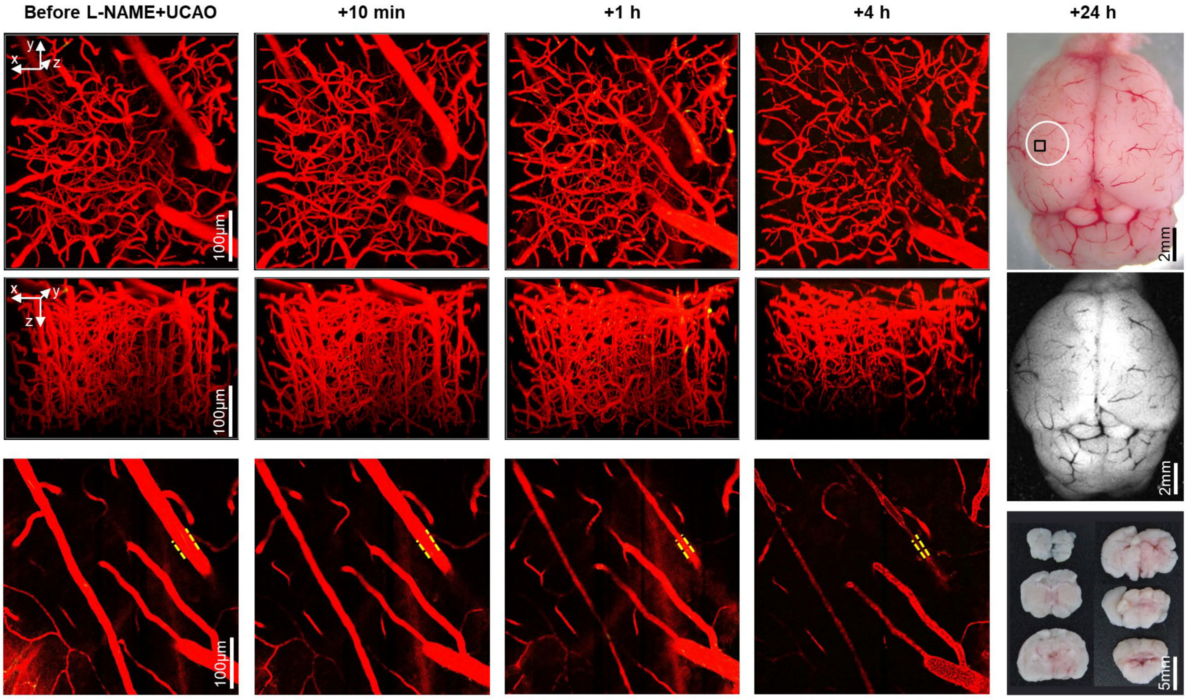


**Supplemental Figure 7. Intravital microscopy shows diffuse arteriolar constriction at 1 and 4 h after L-NAME+UCAO.** Intravital microscopy images and stacks (z-step size=1 μm, total z-depth=300 μm) for the black squared cortical region within the white circle area (top-row, far-right) before vs. 10 min, 1 h, and 4 h after L-NAME+UCAO in a representative BALB/c mouse with cerebral infarction assessed by autofluorescence imaging (middle-row, far-right) and TTC staining (bottom-row, far-right) of the excised brain at 24 h. Texas-red-Dextran-positive arteriolar diameter and vascularity are diffusely decreased; mildly at 10 min, more prominently at 1h, and severely at 4 h (top-row and middle-row). Arteriolar constriction is clearer in the top layer of the stacked images (yellow dashed lines in the bottom-row).

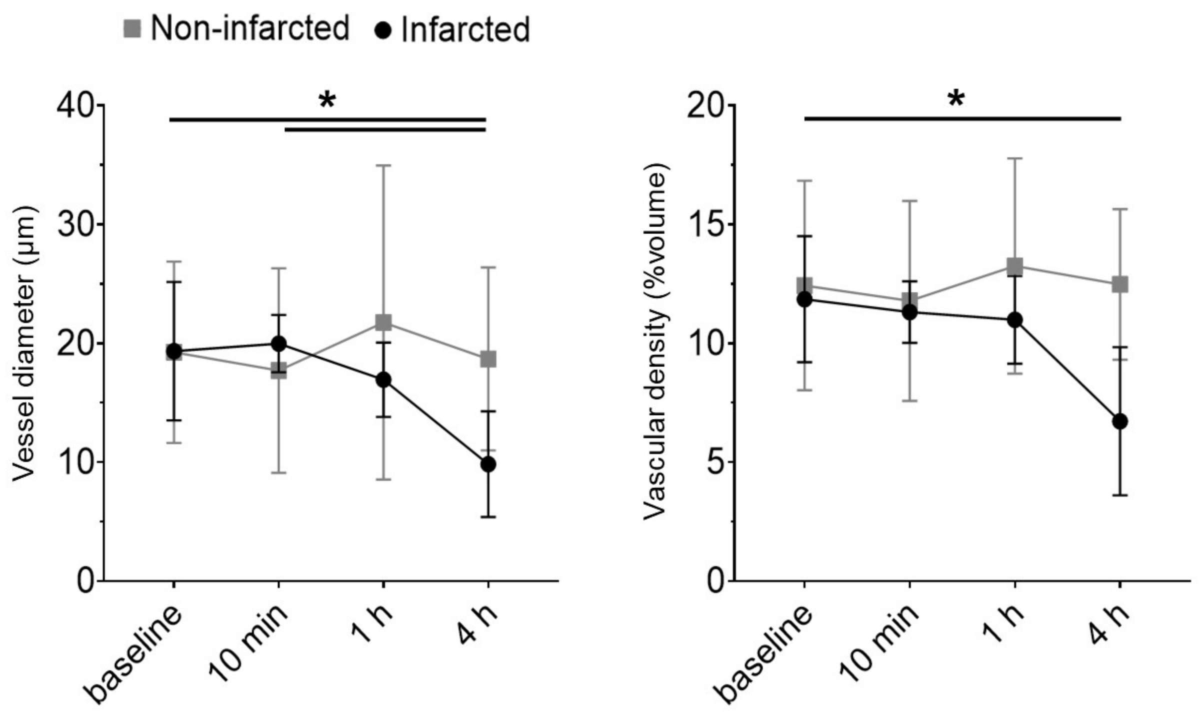


**Supplemental Figure 8. Vascular diameter and density were reduced at 4 h after L-NAME+UCAO in mice with infarction compared to those without.** Changes in cortical vessel diameter (μm) and vascular density (%volume) at 10 min, 1 h, and 4 h after L-NAME+UCAO, compared with the pre-intervention baseline. Infarction occurrence was assessed by TTC staining at ~4 or ~24 h. Graphs represent mean±SE, calculated by quantifying the z-stack data (z-step size=1 μm, total z-depth=300 μm) of intravital microscopy imaging of 21 (three C57BL/6 and 18 BALB/c) mice. $^{*}P<0.05$, linear mixed models with Sidak's multiple comparisons for post-hoc tests.

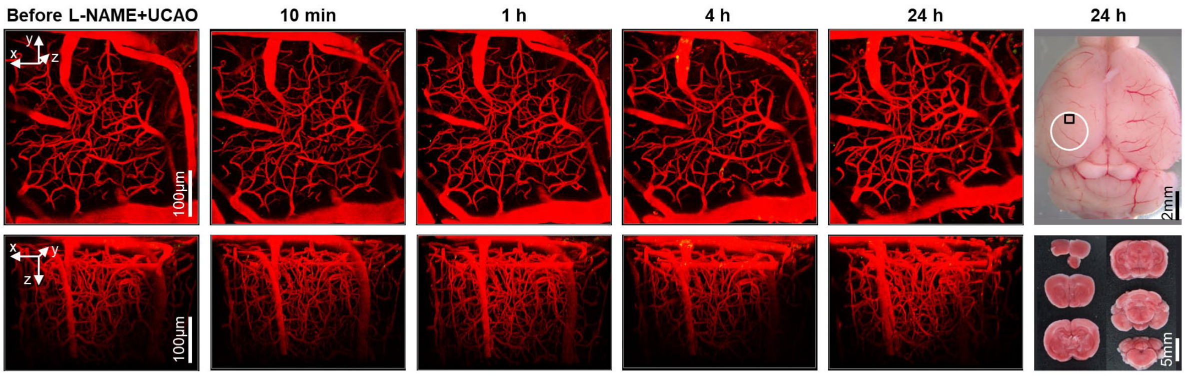


**Supplemental Figure 9. Intravital microscopy shows no significant changes in vascular diameter or density in mice without infarction following L-NAME+UCAO.** Intravital microscopy images and stacks (z-step size=1 μm, total z-depth=300 μm) for the black squared cortical region within the white circle area (top-row, far-right) at baseline and by 24 h after L-NAME+UCAO in a representative BALB/c mouse without cerebral infarction, as assessed by TTC staining performed after the last imaging session.

**Supplemental Video 1. Persistent SDI.** For further details, see Figure 2A in the main text.

**Supplemental Video 2. Transient SDI.** For further details, see Figure 2B in the main text.

**Supplemental Video 3. Transient SDI in the contralateral hemisphere with recovery.** For further details, see Figure 2C in the main text.

**Supplemental Video 4. Intravital microscopy shows real-time constriction and dilatation of cortical arterioles in a BALB/c mouse at ~4 h after L-NAME+UCAO.** Note that blood flow (Texas-red-Dextran with Rhodamine-6G for platelets and leukocytes) becomes sluggish and is then restored in conjunction with vasoconstriction and subsequent vasodilation (arrows). For further details, see Figure 5B in the main text.

**Supplemental Video 5. Intravital microscopy shows real-time rolling and adhesion of Rhodamine-6G-positive platelets and leukocytes on vascular endothelium in a BALB/c mouse at ~4 h after L-NAME+UCAO, unlike at ~1h.**

**Supplemental Video 6. Intravital microscopy shows real-time rolling and adhesion of some Rhodamine-6G-positive platelets and leukocytes on vascular endothelium at ~4 h, but not by 1 h, after L-NAME+UCAO in a BALB/c mouse, followed by firm adherence of numerous cells at ~24 h.**

### Supplemental Results

***Unilateral common carotid occlusion (UCAO) induced cerebral infarction in ~75% of C57BL/6 mice pre-treated with a single intraperitoneal dose of $N_{\omega}$-nitro-L-arginine methyl ester (L-NAME).*** L-NAME+UCAO (n=74 C57BL/6 mice; pre-planned sacrifice at 1 d [n=42], 2 d [n=9], 3 d [n=14], and 6 d [n=9]) induced infarction frequently (Figure 1, A and B).

In the 1 d group, 14% (6/42) died due to large hemispheric infarction (confirmed by 2,3,5-Triphenyltetrazolium chloride [TTC] staining) before the sacrifice time-point (lethal infarction). Mean±standard error (SE) infarct volume of the 36 mice that survived was 115±20 $mm^3$ (median 59, interquartile range [IQR] 0-242); 16 (44%) had large infarcts, 7 (19 %) had small infarcts, and the remaining 13 (36%) did not have an infarct. The overall incidence of infarction including lethal infarction was 69% (29/42). Severe infarction (large infarction>100 $mm^3$ or lethal infarction) occurred in 52% (22/42).

In the 2 d group, lethal infarction occurred in 56% (5/9). Infarct volume of the four mice that survived was 74±30 $mm^3$ (median 52, IQR 34-113); one (25%) had large and the other three (75%) had small infarcts. The overall incidence of infarction including lethal infarction was 100% (9/9), with 66.7% (6/9) severe infarction.

In the 3 d group, lethal infarction occurred in 50% (7/14). Infarct volume of the seven mice that survived was 55±25 $mm^3$ (median 24, IQR 0-119), and two (29%) had large infarcts, two (29%) had small infarcts, and the other three (43%) had no infarct. The overall incidence of infarction including lethal infarction was 79% (11/14), with 64% (9/14) severe infarction.

In the 6 d group, lethal infarction occurred in 56% (5/9). Of the four that survived, one (25%) had a small infarct (36 $mm^3$), and the other three (75%) had no infarct. The overall incidence of infarction including lethal infarction was 67% (6/9), with 64% (5/9) severe infarction.

***L-NAME+UCAO-mediated severe (large or lethal) infarction occurred more frequently in BALB/c than in C57BL/6 mice.*** UCAO alone occasionally induced infarction in BALB/c mice (14%, 4/28; Figure 1A). Twelve of 14 receiving UCAO for 24 h and 12 of 14 receiving UCAO for 7 d had no infarction. Three of the four with infarction had small lesions, whereas the other one (UCAO for 24 h) had lethal infarction. In contrast, L-NAME+UCAO caused infarction far more frequently (76%, 56/74; Figure 1, A and B) by pre-planned sacrifice day 1, 2, 3, or 6. Most of the infarcted mice in this cohort had large lesions (84%, 47/56) when assessed after premature death (91%, 30/33) or sacrifice (74%, 17/23). Infarct volumes of the surviving mice were 126±21, 0, 13±9, and 0 $mm^3$, respectively, and these data also suggest that mice with larger infarcts are more likely to die before sacrifice (Figure 1B). Severe infarction occurred in 71% (30/42), a relatively high incidence compared to that of the C57BL/6 mouse 1 d group (52%, 22/42; $P$=0.07).

In the 1 d group, 31% (13/42) had lethal infarction. Mean±SE infarct volume of the 29 mice that survived was 126±21 $mm^3$ (median 143, IQR 0-206); 17 (59%) had large infarcts, four (14%) had small infarcts, and the other eight (28%) had no infarct. The overall incidence of infarction including lethal infarction was 81% (34/42), with 71% (30/42) severe infarction, which was relatively high compared with the C57BL/6 mouse 1 d group (52%, $P$=0.07).

In the 2 d group, lethal infarction occurred in 67% (6/9). The three animals that survived did not have an infarct.

In the 3 d group, lethal infarction occurred in 64% (9/14). Infarct volume of the five mice that survived was 13±9 $mm^3$ (median 0, IQR 0-26); two (40%) had small infarcts, and the other three had no infarct. The overall incidence of infarction including lethal infarction was 79% (11/14), with 64% (9/14) severe infarction.

In the 6 d group, lethal infarction occurred in 67% (6/9). The three mice that survived did not have an infarct.

***L-NAME+UCAO-mediated infarction occurred less often in SV129 mice***. In SV129 mice, L-NAME+UCAO did induce cerebral infarction, but less frequently (35%, 17/48) than in C57BL/6 and BALB/c (Figure 1, A and B). Most (65%, 11/17) of the infarcted mice in this cohort had large lesions when assessed after premature death (100%, 8/8) or pre-planned sacrifice (33%, 3/9) on day 1, 2, 3, or 6. Infarct volumes of the mice that survived until sacrifice were 26±17, 15±15, 15±12, and 3±2 $mm^3$, respectively. This declining trend again indicates that mice with larger infarcts are more likely to die before the time-points for sacrifice.

In the 1 d group, 17% (2/12) had lethal infarction. Mean±SE infarct volume of the 10 mice that survived was 26±17 $mm^3$ (median 0, IQR 0-9); two mice (20%) had a large infarct, one (10%) had a small infarct, and the other seven (70%) had no infarct. The overall incidence of infarction including lethal infarction was 42% (5/12), with 33% (4/12) severe infarction.

In the 2 d group, 14% (1/7) had lethal infarction. Infarct volume of the six mice that survived was 15±15 $mm^3$ (median 0, IQR 0-0); none (0%) had large cerebral infarcts, and one (17%) had a small infarct (91 $mm^3$), and the other five (83%) had no infarct. The overall incidence of infarction including lethal infarction was 29% (2/7), with 14% (1/7) severe infarction.

In the 3 d group, 15% (2/13) had lethal infarction. Infarct volume of the 11 mice that survived was 15±12 $mm^3$ (median 0, IQR 0-7); one (9%) had a large cerebral infarct, two (18%) had small infarcts, and eight (73%) had no infarct. The overall incidence of infarction including lethal infarction was 39% (5/13), with 23% (3/13) severe infarction.

In the 6 d group, 31% (5/16) had lethal infarction. Infarct volume of the 11 mice that survived was 3±2 $mm^3$ (median 0, IQR 0-0); two (18%) had small infarcts (13 $mm^3$ and 22 $mm^3$), and 9 (85%) had no infarct. The overall incidence of infarction including lethal infarction was 44% (7/16), with 31% (5/16) severe infarction.

Note that subsequent experiments did not include SV129 mice because of infrequent cerebral infarction.

***Administering L-NAME after, rather than before, prolonged UCAO resulted in lower incidence of infarction and smaller lesion size.*** In C57BL/6 mice, L-NAME administration at 3 h, 1, 2, 3, 5, or 7 d after UCAO (n=54, 9/time-point), i.e., UCAO+L-NAME, could also induce infarction. In the 3 h group, 33% (3/9) had lethal infarction before sacrifice at 24 h. Mean±SE infarct volume of the six surviving mice was 135±48 $mm^3$ (159, IQR 0-217); four (67%) had large infarcts, and two (33%) had no infarct. Overall, infarction occurred in 78% (7/9). Unlike in the 3 h group, infarction tended to occur less frequently in the 1, 2, 3, 5, and 7 d groups (vs. the aforementioned 1 d group of C57BL/6 mice with UCAO after L-NAME, i.e., L-NAME+UCAO): no animal died before the 24 h sacrifice time-point (following UCAO for 1~7 d), and infarction occurred in 44% (4/9, two large infarction), 33% (3/9, no large infarction), 44% (4/9, two large infarction), 22% (2/9, one large infarction), and 22% (2/9, two large infarction), respectively. Mean±SE (median [IQR]) infarct volumes of the surviving mice were 54±27 $mm^3$ (0 [0-94]), 2±2 $mm^3$ (0 [0-0]), 49±27 $mm^3$ (0 [0-85]), 34±25 $mm^3$ (0 [0-23]), and 39±26 $mm^3$ (0 [0-35]), respectively.

In BALB/c mice, L-NAME administration 3 h, 1, 2, 3, 5, or 7 d after UCAO (n=54, 9/time-point), i.e., UCAO+L-NAME, could also induce infarction. In the 3 h group, 44% (4/9) had lethal infarction before the 24 h time-point. Infarct volume in the five surviving mice was 151±42 $mm^3$ (median 169, IQR 101-217); four (80%) had large infarcts, and one

(20%) had a small infarct. Overall, cerebral infarction occurred in 100% (9/9). Unlike in the 3 h group, lethal infarction tended to occur less frequently in the 1, 2, 3, 5, and 7 d groups (vs. the aforementioned 1 d group of BALB/c mice with UCAO after L-NAME, i.e., L-NAME+UCAO): lethal infarction and surviving animals' infarction occurred, respectively, in 11% and 63% (1/9 and 5/8), 22% and 86% (2/9 and 6/7), 11% and 88% (1/9 and 7/8), 0% and 44% (0/9 and 4/9), and 0% and 56% (0/9 and 5/9). Among the surviving mice, large infarcts were rarely observed: 13% (1/8), 14% (1/7), 38% (3/8), 11% (1/9), and 0% (0/9), respectively. Mean±SE (median [IQR]) infarct volumes of the surviving mice were 54±27 $mm^3$ (36 [0-64]), 53±21 $mm^3$ (45 [7-79]), 86±36 $mm^3$ (49 [0-152]), 25±14 $mm^3$ (0 [0-48]), and 15±7 $mm^3$ (6 [0-24]), respectively.

***L-NAME+UCAO induced infarction despite cortical blood flow initially being as high as ~65% in the core region.*** The incidence of severe (i.e., large or lethal) infarction was significantly higher in BALB/c mice (56%, 14/25) than in C57BL/6 mice (26%, 5/19; $P$=0.049). The 14 BALB/c mice with severe infarction tended to have lower initial (0-10 min) regional cortical blood flow (rCoBF) (58.9±4.5%) in the core region, compared with the other 10 BALB/c mice (71.4±3.9%; $P$=0.05, Mann-Whitney $U$ test). C57BL/6 mice showed similar results (62.3±4.8% vs. 70.0±2.7%; $P$=0.16, Mann-Whitney $U$ test).

***Spreading ischemia identified mice that progressed to infarction after L-NAME+UCAO.*** Occasionally, spreading ischemia (known to be linked to spreading depolarization [SDI]) spread into the contralateral hemisphere. Of the mice with SDI, 60% (6/10) showed rCoBF recovery (Figure 2, B and C) by the end of the 6 h monitoring period. However, 83% (5/6) mice with rCoBF recovery, as well as 100% (4/4) mice without rCoBF recovery (Figure 2A), had severe infarction by 24 h. These five animals likely had additional SDI bouts without subsequent rCoBF recovery, i.e., terminal or persistent SDI (1), during the remaining (6-24 h) period when laser speckle contrast imaging (LSCI) monitoring was not performed. Additionally, SDI could initiate and spread within the contralateral hemisphere, as shown in a mouse with prior (persistent) SDI in the ipsilateral hemisphere (Figure 2C).

***L-NAME+UCAO-mediated serial changes in core rCoBF differed depending on the occurrence of SDI (with or without rCoBF recovery up to 6 h) and infarction (up to 24 h).*** We stratified all 44 mice (19 C57BL/6 mice and 25 BALB/c mice) that underwent 6 h LSCI monitoring into the following four groups by the occurrence of SDI and cerebral infarction: i) non-infarcted [SDI(-)·Non-infarcted, n=9 C57BL/6 mice and 8 BALB/c mice]; ii) no SDI but infarcted [SDI(-)·Infarcted, n=7 and 9]; iii) SDI with rCoBF recovery but infarcted [SDI(+)·Recovery(+)·Infarcted, n=3 and 3]; and iv) SDI that persisted without rCoBF recovery (until the 6 h time-point) and infarcted [SDI(+)·Recovery(-)·Infarcted, n=0 and 4]. We quantified mean rCoBF for the following four fixed post-L-NAME+UCAO time-periods in all [SDI(+) and SDI(-)] mice: 0-10 and 10-30 min, in which no animals exhibited SDI; a SDI-related period (30-330 min), during which every SDI that occurred was observed in the 6 h monitoring); and, lastly, 330-360 min. We also calculated least squares (LS) mean values for each time-period in each group.

As shown in Supplemental Figure 3A for C57BL/6 mice, initially (at 0-10 min) after L-NAME+UCAO, there were no significant inter-group differences in rCoBF, and the LS mean values were higher than 60% in every group: 63.9% in the SDI(+)·Recovery(+)·Infarcted group and about 70% in the two SDI(-) groups (with or without infarction by 24 h). There was no SDI(+)·Recovery(-)·Infarcted group. During SDI (shaded area in Supplemental Figure 3A), there was a substantial (mean) rCoBF drop to about 30% in the SDI(+)·Recovery(+)·Infarcted group; thereafter (330-360 min), LS mean rCoBF values were about 50%. In the SDI(-) groups, LS mean rCoBF values were about 70% during the 30-330 min and 330-360 min periods. Thus, LS mean rCoBF values during the 30-360 min periods were significantly lower in the SDI(+)·Recovery(+)·Infarcted group than in the SDI(-)·Non-infarcted group.

As shown in Supplemental Figure 3B for BALB/c mice, initially (at 0-10 min) after L-NAME+UCAO, there was a significant difference in rCoBF between SDI(-)·Infarcted and SDI(+)·Recovery(-)·Infarcted group, but the LS mean values were higher than 30% in every group: 44.1% in the SDI(+)·Recovery(-)·Infarcted group, 59.9% in the SDI(+)·Recovery(+)·Infarcted group, and about 70% in the two SDI(-) groups (with or without infarction by 24 h). During SDI (that occurred between the 30-330 min time-points), there was a substantial (mean) rCoBF drop to about 30% in the SDI(+) groups. Post-SDI LS mean rCoBF values were lower than 20% in the SDI(+)·Recovery(-)·Infarcted group, whereas they were about 50% in the SDI(+)·Recovery(+)·Infarcted group. In the SDI(-) groups, LS mean rCoBF values were higher than 65% during the 10-360 min periods. Thus, LS mean rCoBF values during the 10-360 min periods were significantly lower in the SDI(+)·Recovery(-)·Infarcted group than in the SDI(-) groups.

***Serial changes in rCoBF in non-core regions of interest (ROIs) of C57BL/6 and BALB/c mice (n=44) after L-NAME+UCAO, with stratification by the occurrence of SDI (up to 6 h) and infarction (up to 24 h).*** Comparing ROI-3 (ipsilateral anterior cerebral artery territory) with the core region (ROI-1), rCoBF was overall relatively high but with similar patterns of serial changes and inter-group differences in rCoBF (Figure 2D in the main text). In the SDI(+)·Recovery(-)·Infarcted group, post-SDI LS mean rCoBF was as low as ~30%. In the other three groups, LS mean rCoBF values were higher than 50% at all time periods.

ROI-4 (contralateral anterior cerebral artery territory), compared with the ROI-1 and

ROI-3, exhibited similar but less pronounced patterns of serial changes and inter-group differences in rCoBF (Figure 2D in the main text). In addition, post-SDI LS mean rCoBF was significantly lower in the SDI(+)·Recovery(-)·Infarcted group (~60%) than in the other three groups (>80%).

ROI-6 (contralateral region opposite to the ROI-1) had no significant L-NAME+UCAO-mediated rCoBF changes in any groups, except that post-SDI LS mean rCoBF was significantly lower in the SDI(+)·Recovery(-)·Infarcted group (~70%) than in the other three groups (>90%; Figure 2D in the main text).

Finally, analysis of ROI-2 showed mixed ROI-1 and ROI-3 characteristics, while ROI-5 data showed mixed ROI-4 and ROI-6 characteristics (Supplemental Figure 4).

***Infarction following L-NAME+UCAO was not associated with systemic hypotension.*** We stratified 56 mice (27 C57BL/6 and 29 BALB/c mice) by the occurrence of SDI (up to 90 min) and cerebral infarction (up to 24 h). In each group, we calculated LS mean values of each parameter for pre-intervention baseline and the following three fixed post-L-NAME+UCAO time-periods: 0-10 min, in which no animals exhibited SDI; a SDI-related period (10-70 min), during which every SDI that occurred was observed in the 90 min monitoring; and, lastly, 70-90 min. In both SDI(-)·Non-infarcted group (n=19, 10 C57BL/6 and 9 BALB/c mice) and SDI(+)·Infarcted group (n=5, 1 C57BL/6 and 4 BALB/c mice), systolic and diastolic BP significantly increased (up to ~30 mmHg and ~20 mmHg, respectively) after L-NAME+UCAO, without significant inter-group differences in any time periods (Figure 3E). In the SDI(+)·Infarcted group, there were no significant SDI-related BP changes (shaded area).

In the saline control group (n=5 C57BL/6 and 5 BALB/c mice; Figure 3B in the main text), no animals had an SDI or an infarct. When compared with the pre-intervention baseline, heart rate and systolic BP were slightly higher at 70-90 min, whereas diastolic BP was slightly lower at 10-70 and 70-90 min. There was no significant serial change in rCoBF in the core region (ROI-1).

In the L-NAME only group (n=5 C57BL/6 and 5 BALB/c mice; Figure 3C in the main text), no animal had either an SDI or an infarct. Heart rate was slightly lower at 10-70 min after L-NAME administration (vs. baseline). As expected, both systolic and diastolic BPs elevated significantly (~10 mmHg) higher at 0-10 min, compared with baseline. Systolic and diastolic BPs rose further (~10 mmHg) at 10-70 min, reaching a plateau. Unlike in BP, rCoBF did not change significantly.

In contrast to the previous two groups, in the UCAO only group (n=5 C57BL/6 and 5 BALB/c mice; Figure 3D in the main text), one BALB/c mouse had an SDI at ~40 min, with acute rCoBF drop (from 30% to 10%) and partial recovery (from ~30% to ~10% and then to ~20%) in the core region (upper graph in the shaded areas of Supplemental Figure 6A). Heart rates at 10-70 and 70-90 min after UCAO (vs. baseline) were slightly higher with an increasing trend, whereas both systolic and diastolic BPs at 0-10, 10-70, and 70-90 min (vs. baseline) were slightly lower with a decreasing trend (Supplemental Figure 6A). At 24 h, this animal did not have an infarct. The other nine mice without SDI (Figure 3D in the main text) exhibited no significant serial changes in systolic or diastolic BP after UCAO, although heart rates were significantly (~50/min) higher at 10-70 and 70-90 min with an increasing trend. As expected, UCAO significantly decreased rCoBF in the core region, to ~60% of the baseline value.

The L-NAME+UCAO group (n=12 C57BL/6 and 14 BALB/c mice) had 19 SDI(-)·Non-infarcted mice (10 C57BL/6 and nine BALB/c; Figure 3E in the main text), two SDI(-)·infarcted mice (one C57BL/6 and one BALB/c; Supplemental Figure 6B), and five SDI(+)·Infarcted mice (one C57BL/6 and four BALB/c; Figure 3E in the main text). In the

SDI(-)·Non-infarcted group, heart rates were slightly higher at 0-10 and 70-90 min (vs. baseline), which was not the case in the SDI(+)·Infarcted group; however, heart rates had no significant inter-group differences in any time period. Both the SDI(-)·Non-infarcted group and the SDI(+)·Infarcted group showed significantly elevated systolic and diastolic BP (up to ~30 mmHg and ~20 mmHg, respectively) after L-NAME+UCAO, without notable inter-group differences in any time period. Moreover, there were no significant SDI-related BP changes within the SDI(+)·Infarcted group (see the shaded area of the BP graph in Figure 3E in the main text). In line with the aforementioned LSCI experiments that did not involve monitoring heart rate or BP (Figure 2D in the main text), L-NMAE+UCAO-mediated initial reduction of rCoBF in ROI-1 was significantly larger in the SDI(+)·Infarcted group than in the SDI(-)·Non-infarcted group, although LS mean rCoBF values were again higher than 30% in both groups (~60% and ~40%, respectively). Moreover, further SDI-related reduction in rCoBF (to below 30%) was observed in the SDI(+)·Infarcted group, while rCoBF remained at ~60% in the SDI(-)·Non-infarcted group. Serial heart rate, BP, and rCoBF data for the two SDI(-)·Infarcted mice (Supplemental Figure 6B) were similar to those of the SDI(-)·Non-infarcted group.

Lastly, none of the monitoring showed significant inter-strain differences, though control group heart rates were faster in C57BL/6 mice than in BALB/c mice, in all time periods (Data not shown). To summarize, combined monitoring of heart rate, BP, and cerebral perfusion indicated that L-NAME+UCAO-mediated, SDI-related induction of ischemic stroke is not due to systemic hypotension.

***Streptozotocin (STZ)- and high-fat diet (HFD)-mediated hyperglycemia and hyperlipidemia in C57BL/6 and ApoE$^{-/-}$ mice.*** As shown in Figure 6A, fasting glucose levels were significantly higher in STZ and HFD+STZ groups than in non-treated groups (~400 mg/dL and ~200 mg/dL, respectively) at 7 d before UCAO in each strain (all $P<0.001$). Total cholesterol levels (which were significantly higher in ApoE$^{-/-}$ mice than in C57BL/6 mice, as expected) were ~two-fold higher in the HFD+STZ group than in either saline or HFD group in each strain. In ApoE$^{-/-}$ mice, the STZ and HFD+STZ group had similarly high levels of total cholesterol. Triglyceride levels were notably high in the HFD+STZ groups in both strains, particularly in C57BL/6 mice (Mean±SE, 1692±311 mg/dL vs. 744±139 mg/dL, $P=0.007$), compared with the other three (saline, STZ, and HFD) group (~200 mg/dL, all $P<0.001$). High-density lipoprotein levels were significantly elevated in C57BL/6 mice (>~60 mg/dL) compared to ApoE$^{-/-}$ mice (<~40 mg/dL, all $P<0.001$). In contrast, low-density lipoprotein levels were significantly higher in the STZ or HFD group of ApoE$^{-/-}$ mice (>~150 mg/dL) than in the saline, STZ, or HFD group of C57BL/6 mice (~10 mg/dL). However, the HFD+STZ group of C57BL/6 mice had increased low-density lipoprotein levels (94±14 mg/dL) compared to the saline group of ApoE$^{-/-}$ mice (63±5 mg/dL, $P=0.042$).

### Supplemental Methods

#### *Study design for preclinical research*

***Experiment 5.*** High-resolution microCT-based thrombus imaging and synthesis of fibrin-targeted gold nanoparticles were performed, as we previously published (2, 3). Here, three C57BL/6 mice were additionally used to verify that the *in vivo* direct thrombus imaging technique can clearly visualize and monitor cerebral thromboembolism before and after tissue plasminogen activator therapy (25 mg/kg, 600 μL) in an embolic stroke model (4, 5). For intravital two-photon microscopy imaging (IVIM Technology, Daejeon, Korea) of blood flow and white blood cells / platelets, Texas red Dextran (Thermo Fisher Scientific, Boston, MA, USA) and Rhodamine 6G (TCI, Tokyo, Japan) were used, as we previously reported (6, 7).

Data from 16 mice that received L-NAME + left UCAO were used for the following quantitative analysis, after exclusion of 7 mice due to: anesthesia failure (n=1 C57BL/6 mice), poor quality of the images (n=1 C57BL/6 and 2 BALB/c mice), and death <1 h after L-NAME+UCAO (n=3 BALB/c mice). Quantification of z-stack images (z-step size=1 μm, total z-depth=300 μm) of the 16 (three C57BL/6 and 13 BALB/c) mice was performed using FIJI ImageJ v1.54f (National Institutes of Health, Bethesda, MD, USA). In brief, z-stack images were down-sampled to 256×256 pixels, and a Gaussian 3D filter (2×2×2 pixels) was applied. After the images were binarized, vascular density was calculated as % volume of all segmented vessels. Then, mean vessel diameter (μm) was calculated using two FIJI plugins (‘Skeleton 2D/3D’ and ‘Local thickness’). To average the local thickness values along the vessel skeleton, non-visualized parts of the vessel were approximated using the pre-L-NAME+UCAO baseline data as a reference, and their thicknesses were set to 0. For histology, brains were harvested following cardiac perfusion. After fixing 2-mm thick coronal sections in a formaldehyde solution for ~24 h, 4-μm thick sections containing middle cerebral arteries were carefully prepared (3) using a microtome (LEICA [RM2235], Nussloch, Germany) for H&E staining (Abcam, Cambridge, UK). Representative image files were captured (n=3/mice) using a microscope (Olympus [DP73], Tokyo, Japan) with cellSens imaging software (Olympus, Tokyo, Japan).

***Experiment** 7.* A total of 168 (n=92 C57BL/6 and 76 ApoE$^{-/-}$) 11-week-old mice were randomly divided into the following groups: vehicle control, STZ-treated, HFD-fed, and HFD+STZ-treated groups. STZ was administered via intraperitoneal injections at 50 mg/kg/day (Sigma, St. Louis, MO, USA) in 50 mM citrate buffer for 5 consecutive days. The non-STZ groups received the citrate buffer vehicle. HFD consisting of 60% fat, 20% carbohydrates, and 20% protein (Research Diets, Inc., New Brunswick, NJ, USA) lasted for

20 d (8). The two non-HFD groups were fed a normal chow diet. Whole blood glucose levels were measured by using a blood glucose meter (ACCU-CHEK; Roche, Indianapolis, IN, USA) following a 6 h fasting, 7 d after the last administration of STZ or vehicle. UCAO was performed 7 days later. Serum cholesterol levels were measured 24 hours later using the Clinical Chemistry Analyzer AU480 (Beckman Coulter, Brea, CA, USA) and SEKURE® Clinical Chemistry Kits (Sekisui, Tokyo, Japan). Next, TTC staining of the brain was performed. A total of 134 (n=75 C57BL/6 and 59 $ApoE^{-/-}$) mice were included in the final analysis after 34 mice were excluded for the following reasons: STZ treatment-related death (n=19), absence of predefined hyperglycemia (<250 mg/dL) (9) at 7 d after STZ treatment (n=4), and poor TTC staining for reliably assessing infarct presence (n=11). In 20 (10 HFD+STZ and 10 vehicle control) of the C57BL/6 mice (without UCAO), asymmetric dimethylarginine (ADMA) and symmetric dimethylarginine (SDMA) concentrations were also measured by (QuBEST BIO, Yongin, Korea) using a liquid chromatography-mass spectrometry method with Agilent 6490 triple quadrupole spectrometer (Agilent Technologies, Basel, Switzerland)

***Analysis of LSCI data.*** In Experiment 1, LSCI flux values of mice in prone position can be altered by transitioning from prone to supine to ensure a secure UCAO before returning to the prone position to resume monitoring. Thus, we adjusted rCoBF values of every ROI in each animal through two steps: i) setting the initial (0-10 min) mean value at ROI-6 to 100% and ii) adding the difference (100 minus pre-normalization value at ROI-6) to the values of the other five ROIs in order to generate adjusted rCoBF values. This adjustment was not required for Experiment 4 (see the main text), in which mice were kept prone to securely measure BP using an intraarterial catheter, and UCAO was blindly performed without direct manipulation of the ligation site. The procedure involved tightening a loosened knot prepared before LSCI monitoring.
